# Metabolic adaptations in a range expanding arthropod

**DOI:** 10.1101/043208

**Authors:** Katrien H. P. Van Petegem, David Renault, Robby Stoks, Dries Bonte

**Affiliations:** Department of Biology, Ghent University, Ghent, Belgium; UMR CNRS 6553 Ecobio, Université de Rennes 1, Rennes Cedex, France; Evolution and Conservation, KU Leuven, Leuven, Belgium

**Keywords:** essential amino acids, common garden, life-history evolution, GC-MS metabolomics, global change, *Tetranychus urticae*

## Abstract

Despite an increasing number of studies documenting life-history evolution during range expansions or shifts, we lack a mechanistic understanding of the underlying physiological processes. In this explorative study, we used a metabolomics approach to study physiological changes associated with the recent range expansion of the two-spotted spider mite (*Tetranychus urticae*). Mite populations were sampled along a latitudinal gradient from range core to edge and reared under benign common garden conditions for two generations. Using Gas Chromatography-Mass Spectrometry (GC-MS), we obtained metabolic population profiles, which showed a gradual differentiation along the latitudinal gradient, indicating (epi)genetic changes in the metabolome in association with range expansion. These changes seemed not related with shifts in the mites’ energetic metabolism, but rather with differential use of amino acids. Particularly, more dispersive northern populations showed lowered concentrations of several essential and non-essential amino acids, suggesting a potential downregulation of metabolic pathways associated with protein synthesis.

## Introduction

During range expansions or range shifts, species’ life histories can evolve on ecological timescales (Phillips, Brown & Shine 2010). Changing environmental conditions force species to locally adapt and spatial assortment of dispersive phenotypes leads to increased dispersiveness at the expanding/shifting range edge (Shine, Brown & Phillips 2011). These evolutionary processes of local adaptation and spatial selection affect key life-history traits like fecundity, development and dispersal (reviewed in Chuang & Peterson 2015). We therefore expect range edge populations to exhibit physiological adaptations that underlie these observed trait evolutions. Such adaptations should be especially significant in energy-producing pathways, and more particularly in glycolysis (Eanes 2011). Indeed, any elevation of the performance of one life-history trait augments its energetic and metabolite demands at the expense of other traits (*cfr*. the “Y” model of resource allocation, Van Noordwijk & de Jong 1986; Zera & Harshman 2001), thus modifying the global metabolic network operation. For instance, variations in lipid biosynthesis in association with dispersal strategy highly impact metabolite fluxes through lipid pathways (see Zera 2011 for a review). As a result, life-history differentiation is expected to be associated with changes in the metabolome (*i.e.* the set of circulating metabolites within an organism, Oliver *et al.* 1998), as was for example found for ageing in *Caenorhabditis elegans* (Fuchs *et al.* 2010) and reproduction in the Malaria Mosquito *Anopheles gambiae* (Fuchs *et al.* 2014).

Metabolomics is a convenient technique which can be used as a candidate approach to explore an organism’s response to environmental variations and forthcoming environmental changes (Hines *et al.* 2007; Miller 2007; Viant 2008; Bundy, Davey & Viant 2009; Lankadurai, Nagato & Simpson 2013; Hidalgo *et al.* 2014). It provides information on the interaction between an organism’s physiology and its natural environment by identifying metabolites of low to moderate molecular mass within the whole body, cells, tissues or biofluids. Compared to other -omics technologies like genomics and transcriptomics, metabolomics has the significant advantage to focus on ‘downstream’ cellular functions (Snart, Hardy & Barrett 2015), providing a more direct picture of the functional links between causes and consequences of environmental variation (Foucreau *et al.* 2012). Essentially, metabolomics can potentially provide a link between genotypes and phenotypes (Fiehn 2002). When applied on individuals originating from different localities from range core to edge, but reared for several generations under common garden conditions, it should provide insights on the physiological adaptations that underlie life-history evolution during range expansion.

Though a consideration of the whole-organism physiology allows a better understanding of how life-history evolution in natural populations might occur and why this evolution is sometimes constrained (Zera *et al.* 2001;Zera 2011; Ricklefs & Wikelski 2002), few studies documented metabolic variation in wild populations along natural gradients (Sardans, Penuelas & Rivas-Ubach 2011). Instead, most studies assess plastic or evolutionary responses to environmental stressors by manipulating abiotic variables in controlled environments (Sardans *et al.* 2011; Colinet *et al.* 2012; Padfield *et al.* 2016). Notable exceptions are the studies on *Arabidopsis lyrata* that demonstrated distinct metabolic phenotypes along the species’ latitudinal distribution, with a typical cold-induced metabolome in the north, indicating adaptation to the local climate (Davey *et al.* 2008; Davey, Woodward & Quick 2009), but no difference in metabolic fingerprint between large connected *versus* marginal fragmented populations (Kunin *et al.* 2009). These studies, however, used plants that were grown from seeds collected directly from the field. Environmental maternal effects can therefore not be excluded.

The two-spotted spider mite, *Tetranychus urticae* Koch (Acari, Tetranychidae; Fig. 1), a generalist pest species in greenhouses and orchards, expanded its European range from the Mediterranean to at least southern Scandinavia (K. H. P. Van Petegem, personal observation in 2011) during the last decades (for more information, see Carbonnelle *et al.* 2007). Previous research with *T. urticae* showed quantitative genetic life-history differentiation along this latitudinal gradient, with daily fecundity, lifetime fecundity and longevity decreasing from range core to edge, and egg survival, dispersal propensity and sex ratio increasing from range core to edge (Van Petegem *et al.* 2016). We expected this life-history differentiation to be associated with an evolution of the species’ intermediary metabolism, which would manifest into distinct metabolic phenotypes among populations sampled along its expansion gradient.

**Figure 1.**
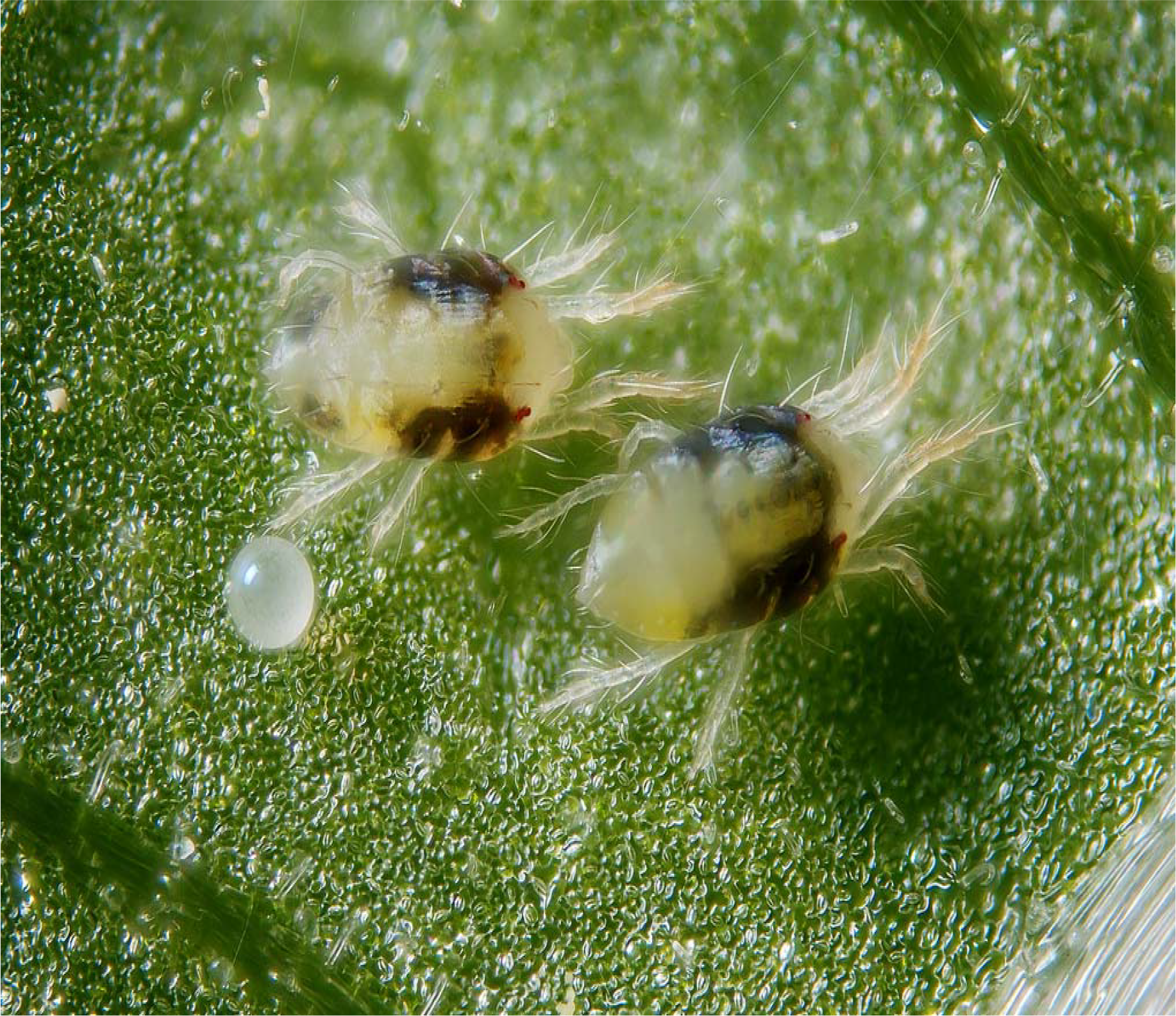
Picture showing two females and one egg of the two-spotted spider mites (*Tetranychus urticae* Koch; Acari, Tetranychidae).

Using a metabolomics approach, the current study aimed to test (i) whether the metabolome of *T. urticae* evolved during its recent range expansion; *i.e.* whether a gradual change in the mite’s intermediary metabolism is present from range core to range edge, showing in the appearance of progressively distinct metabolic phenotypes (metabotypes) (ii) whether this metabolic differentiation could be associated with the up-or downregulation of certain metabolic pathways, for instance enhanced glycolytic activities or lipid metabolism and (iii) whether this evolutionary change in the species’ metabotype is associated with the life-history differentiation that has occurred during its range expansion.

## Materials and methods

### Field sampling and common garden

In August 2012, we hand-sampled mites from nine localities (one population per locality) along an 800 km latitudinal gradient from northwestern Belgium to northern Denmark (Fig. 2). Mites were found on infested leaves of *Lonicera periclymenum* (European honeysuckle, five populations) at high latitudes and on *Euonymus europaeus* (European spindle, one population), *Humulus lupulus* (common hop, one population),–*Sambucus nigra* (European black elderberry, one population) or *Lonicera periclymenum* (European honeysuckle, one population) at lower latitudes. (More information is provided in appendix A.1.) In the laboratory, fifty to several hundreds of mites per population were put on separate whole bean plants (*Phaseolus vulgaris*, variety Prélude -a highly suitable host for *T. urticae*, see Agrawal et al. 2002; Gotoh et al. 2004) and kept under controlled conditions at room temperature with a light-regime of 16:8 LD. After one generation, ten adult female mites per population were taken from their bean plant and put on a piece of bean leaf (±30cm^2^) on wet cotton in a Petri dish. Two such Petri dishes were prepared for each population. The Petri dishes were then used to create a pool of synchronised two-day adult female mites for each population (two-day adult females were preferred, since these are significantly bigger than fresh adults). For this purpose, all females were allowed to lay eggs during 24 hours in a climate room at 27 °C (an optimal temperature for our study species, see Sabelis 1981), with a light-regime of 16:8 LD. The resulting same-aged eggs were subsequently left to develop until they were two-day adult mites, of which only females (which are easily visually recognised) were selected. As mites were kept in common garden for two generations, all direct environmental effects were excluded. Furthermore, since the common garden conditions were optimal for *T. urticae* (bean as host plant, 27°C, 16:8 LD and relatively low mite densities), we could reasonably assume that potential differences among metabotypes of mites from different origins (populations) did not result from differential responses to restrictive/stressful rearing.

**Figure 2.**
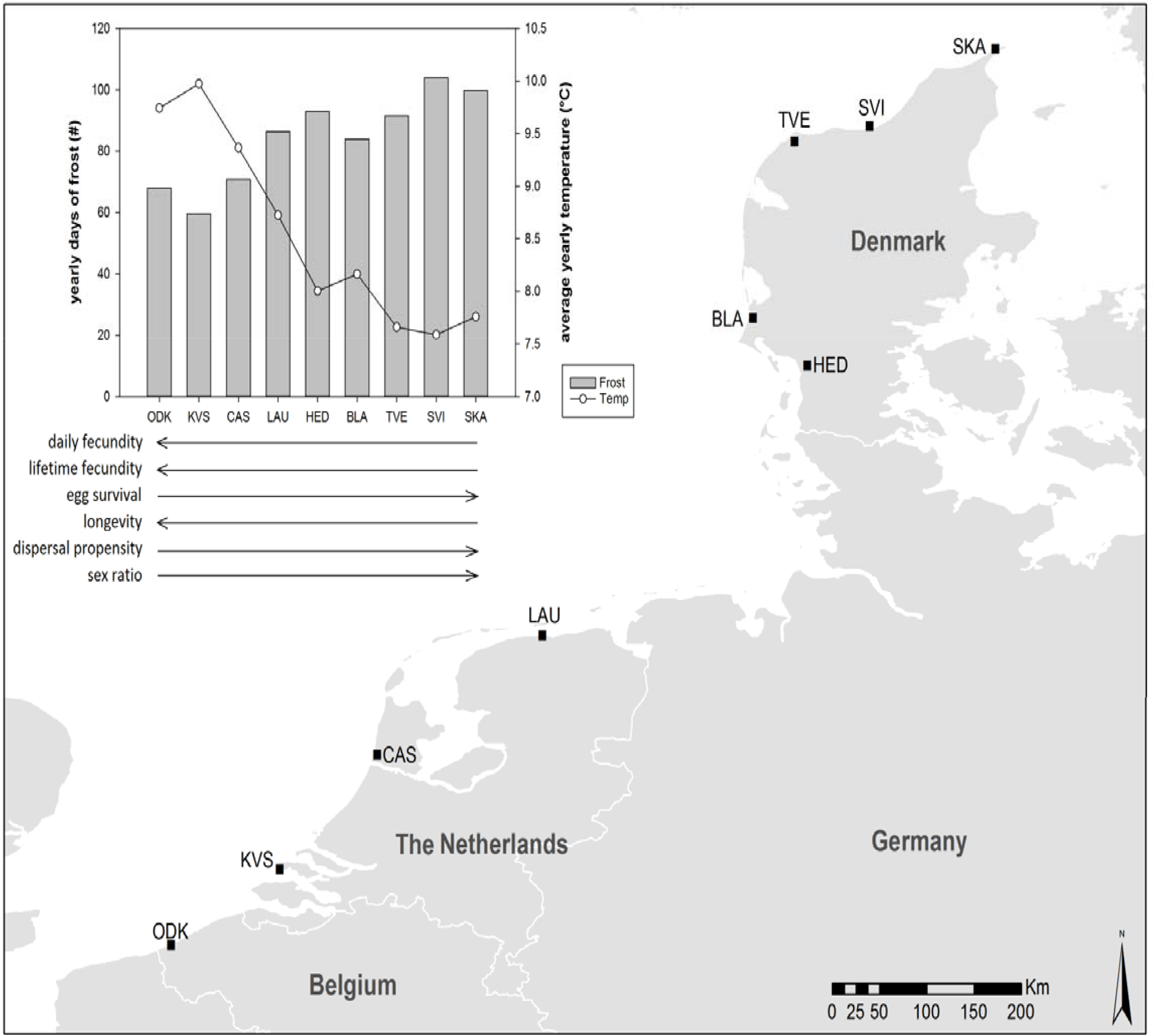
The map shows the nine field collection sites, which are situated in Belgium, The Netherlands and Denmark. The graph shows the yearly number of frost days and the average yearly temperature for each collection site along the latitudinal gradient. These climatic data were obtained from FetchClimate (Microsoft Research, Cambridge) and were averaged over a period of 35 years (1980 to 2015). Below the graph, arrows for each of six life-history traits depict their trend along the latitudinal gradient (increase, decrease). (For more detailed information, see appendix A.l and Van Petegem *et al.* 2015.)

### Metabolomic profiling using Gas Chromatography-Mass Spectrometry (GC-MS)

As we wanted to scan metabolites from different metabolite families (because of their various but connected roles in general organismal physiology), we used GC-MS metabolomics (Koek *et al.* 2011; Khodayari *et al.* 2013). For each population, we constructed the metabolomic profile of five replicated pooled sets of fifty two-day-adult female mites. Each set was placed in a microtube and directly transferred to −80 °C. To be able to measure true quantities of metabolites, it is important to standardise the initial masses of each extract. However, even when pooling fifty individuals, the masses of the replicates were too low to be accurately measured (the measurement error for the mass of the microtube was greater than the summed mass of the fifty mites). Yet, previous research showed that female adult size does not differ among the nine sampled populations (Van Petegem *et al.* 2016). We could thus confidently use and interpret metabolite concentrations in nmol/sample. The samples were first homogenised in ice-cold (−20 °C) methanol-chloroform (2:1), using a tungsten-bead beating equipment (RetschTM MM301, Retsch GmbH, Haan, Germany) at 25 Hz for 1.5 min. After addition of ice-cold ultrapure water, the samples were centrifuged at 4,000 g for 5 min at 4 °C. The upper aqueous phase was then transferred to new chromatographic glass vials, dried-out and resuspended in 30 μl of 20 mg L-l methoxyamine hydrochloride (Sigma-Aldrich, St. Louis, MO, USA) in pyridine and incubated under automatic orbital shaking at 40 °C for 60 min. Subsequently, 30 μl of N-methyl-N-(trimethy Isily I) trifluoroacetamide (MSTFA; Sigma, #394866) was added and the derivatisation was conducted at 40 °C for 60 min under agitation. The samples were then analysed in a GC-MS system (Thermo Fischer Scientific Inc., Waltham, MA, USA), using the same settings as in Khodayari *et al.* (2013). For this purpose, one microliter of each sample was injected in the GC-MS system using the split mode (split ratio: 25:1). After that, the selective ion monitoring (SIM) mode (electron energy: −70 eV) was used to search for the sixty primary metabolites that are most often found in arthropod samples and that were included in our spectral database (see appendix A.2 for a complete overview of these sixty metabolites). The SIM mode ensured a precise annotation of the detected peaks. The calibration curves were set using samples consisting of sixty pure reference compounds at concentrations of 1, 2, 5, 10, 20, 50, 100, 200, 500, 750, 1000, 1500 and 2000 μM. Chromatograms were deconvoluted using XCalibur v2.0.7 software (Thermo Fischer Scientific Inc., Waltham, MA, USA). Finally, metabolite concentrations were quantified according to their calibration curves.

### Statistics

A total of forty-three metabolites were identified.

In a first step, we examined whether distinct metabotypes existed among the sampled populations of *T. urticae* by running a MANOVA. Nine metabolites were first removed from the dataset because they showed a more than 85% correlation with (an)other metabolite(s). Then, all remaining metabolites were log-transformed to obtain a normal distribution, which allowed to proceed to the MANOVA.

In a second step, we tested whether the metabolic profile was gradually changing as a function of latitude (from range core to edge), or as a function of one of the six life-history traits that were previously shown to covary with latitude (daily and lifetime fecundity, egg survival, longevity, dispersal propensity and sex ratio; see Van Petegem *et al.* 2016, see also appendix A.3). The concentrations of all forty-three metabolites were first auto-scaled and transformed (the transformation that best fitted and normalised the data was retained: cube root when looking for covariation with daily fecundity and egg survival; log for latitude; no transformation for lifetime fecundity, longevity, dispersal propensity and sex ratio). Then, metabolic differences among the nine populations were visualised using Partial Least – Discriminant Analysis (PLS-DA). This multivariate analysis was performed using MetaboAnalyst 3.0 (Xia *et al.* 2009; Xia *et al.* 2012; Xia *et al.* 2015). By ordering the populations according to their latitude or according to one of the six life-history traits covarying with latitude, it was possible to check for trends in the metabolite concentrations. To validate the significance of this interpopulation variation, permutation tests (2000 permutations) were run for replicates (with 5 replicates per population) using separation distance (B/W) test statistics. The PLS-DA provided Variable Importance in Projection (VIP) scores, which gave a first overview of the possible existence of a general pattern in the concentrations of quantified metabolites along our invasion gradient: low VIP-scores depict a weak and high scores a strong global pattern. Using a step-wise procedure, only those metabolites with a VIP score of at least 1.2 (1.0 for egg survival because removing the metabolites with a score between 1.0 and 1.2 resulted in a decreased percentage of variation explained) for the first and/or second component were retained for further analysis (compared to 0.8 in Tenenhaus 1998).

In a third step, univariate analyses were performed to test, metabolite by metabolite, whether the global patterns obtained in the previous step could be confirmed. As we aimed at determining if the metabolite levels showed latitudinal patterns along the expansion, we did not run an ANOVA on individual metabolites, but rather processed regressions. Using SAS 9.4 (SAS Institute Inc. 2013), the linear regressions were run for all influential metabolites (VIP scores >1.2, except for egg survival, as mentioned above). Because all five collected replicates per population originated from only one field sample, we ran the regressions using population averages. As our study is explorative, we wanted to avoid false negatives (with the chance of making a Type II error). We therefore did not correct for multiple comparisons (*e.g.* Bonferroni correction). Given the large number of statistical tests, the limited number of populations, and the fact that we highly smoothed differences among populations by rearing the specimens under common garden conditions for two generations (which is atypical for metabolomics studies, where organisms are usually subject to a stressor to elicit a strong response), such a correction would have greatly diminished the statistical power.

A final step linked the selected individual metabolites with one or more metabolic pathways, thus identifying those pathways that were potentially up-or downregulated during the range expansion of *T. urticae*. Pathway enrichment analyses were performed in Metaboanalyst 3.0 (Xia *et al.* 2009; Xia *et al.* 2012; Xia *et al.* 2015) with those metabolites that showed significant effects in the univariate analyses of step three. These pathway analyses were performed with a Fisher’s exact test algorithm, which we ran using the metabolic pathways of *Drosophila melanogaster* (no closer relative of *T. urticae* was available in the program, but primary metabolites are anyway highly conserved, especially among non-blood feeding arthropods). The algorithm calculates the match (number of hits) between the metabolites in a dataset and the totality of metabolites present in a specific pathway. Furthermore, it uses a pathway topology analysis to compute a value for the impact of these metabolites on the pathway. As multiple comparisons are made, corrected Holm p-values are provided.

## Results

### MANOVA

The metabotypes of the female mites significantly differed among the nine sampled populations (F_184,160_=2.2, p<0.001). In further analyses, the potential covariation of metabolite levels with latitude or one of the life-history traits covarying with latitude was then assessed.

### Latitudinal covariation

The PLS-DA showed a separation among the nine populations, which was visible on 3D score plots (see appendix A.4). Of the forty-three identified metabolites, seventeen had VIP scores of at least 1.2 and were thus retained for further analysis (Fig. 3A). They showed a clear general trend from high values in southern to low values in northern populations (Fig. 3A).

**Figure 3.**
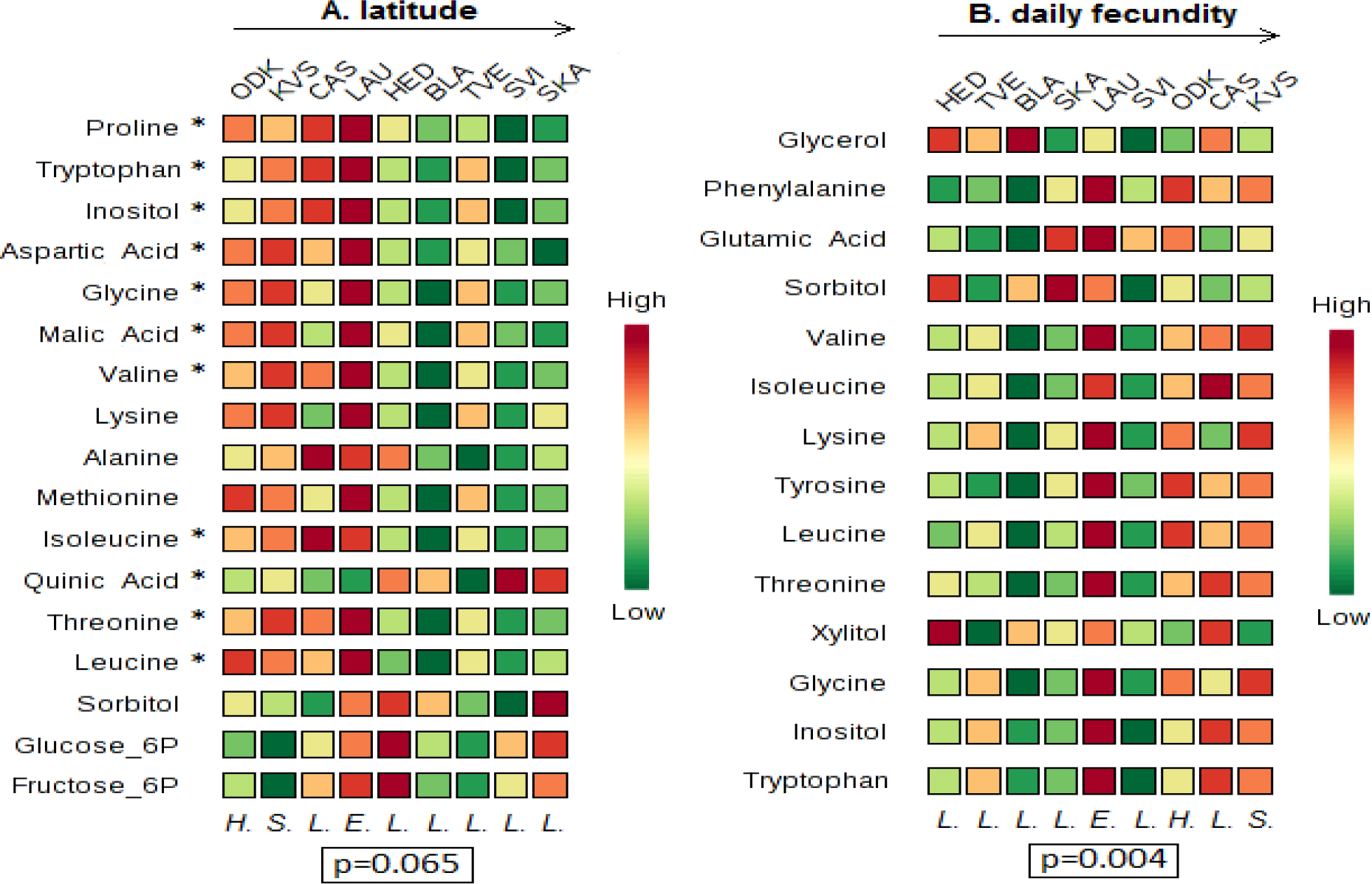
Variable importance plots resulting from the multivariate analyses (PLS-DA) on the metabolomic data. These plots list those metabolites that, based on their VIP score, contribute the most to explaining the variation among the nine populations in our dataset (ODK, KVS, CAS, LAU, HED, BLA, TVE; SVI, SKA). The metabolites are ordered from high to low VIP scores for component 1 (an overview off all scores for component 1 and 2 is provided in appendix A.5). The colour codes indicate the relative concentration of a given metabolic compound for each population (green=low concentration, to red= high concentration). The populations themselves are ordered according to their latitude (A), or from low values at the left to high values at the right for a given life-history trait (daily (B) or lifetime (C) fecundity, egg survival (D), longevity (E), dispersal propensity (F) or sex ratio (G)). For example, in Fig. 3A, LAU is the population with the highest concentration of proline and ODK is the southernmost population (lowest latitude). Below each column (population), a letter signifies the host plant species from which mites were sampled in this population *H.= H. lupulus, S.= S. nigra, L.= L. periclymenum, E.= E. europaeus*). At the bottom of each plot, the p-value resulting from the permutation test is given. At the left side of each plot, an asterisk next to a metabolite name indicates a significant correlation between this metabolite and latitude or the denoted life-history trait. For example, in Fig. 3A, proline shows a significant negative correlation with latitude. (A detailed overview, including p-and F-values, of all linear regressions is found in appendix A.6.)

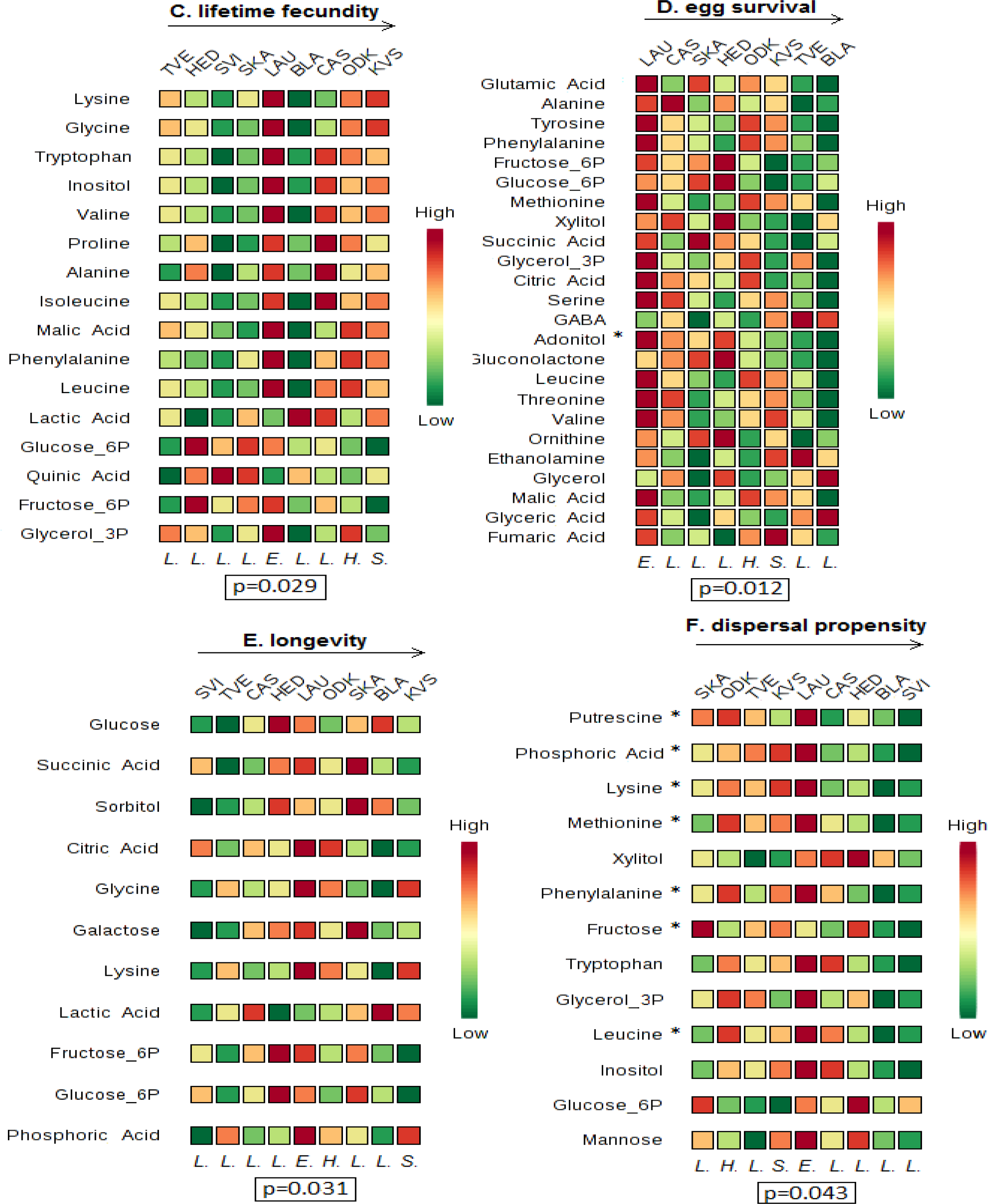

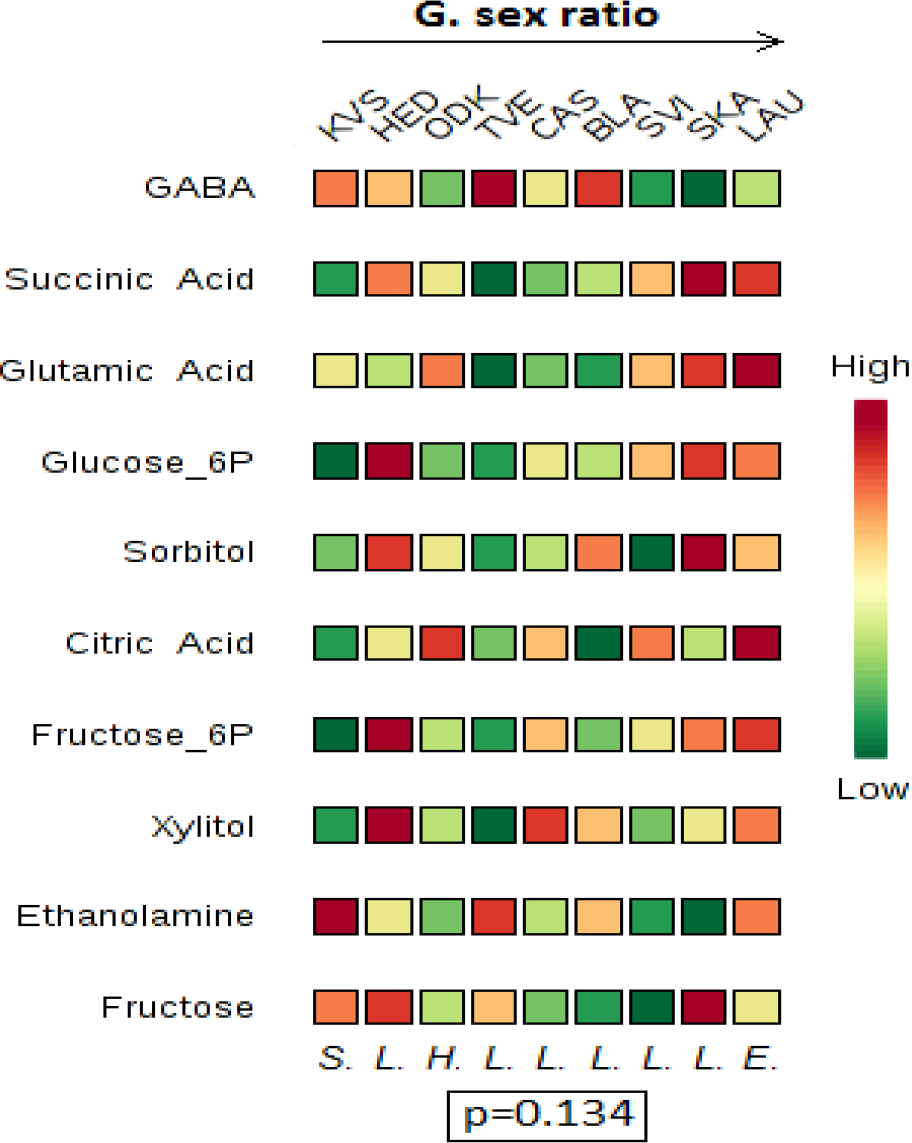

In the subsequent linear regressions, eleven of these seventeen metabolite concentrations varied significantly: ten decreased and one increased with increasing latitude (Fig. 3A and appendix A.6). Among these eleven metabolites, five essential amino acids, three non-essential amino acids (see Rodriguez & Hampton (1966) for an overview of all essential amino acids in *T. urticae* -we defined tryptophan, which is not included in this overview, as essential) and one intermediate of the citric acid cycle can be mentioned.

Pathway analysis indicated that of these eleven metabolites, eight play a significant role in the aminoacyl-tRNA biosynthesis (total: 67, hits: 8, impact=0, Holm p=2.7062E-6) and four in the valine, leucine and isoleucine biosynthesis (total: 13, hits: 4, impact=0.9999, Holm p=4.0129E-4) (both pathway maps are provided in appendix A.7).

### Life history covariation

The PLS-DA showed a separation between the nine (eight for egg survival, for which no data were available for population SVI) populations, which was visible on 3-D score plots (see appendix A.4). Of the forty-three identified metabolites, only those which explained most of the interpopulation variation for a certain life-history trait (high VIP score) were retained for further analysis. Fourteen were retained for daily and sixteen for lifetime fecundity, twenty were retained for egg survival, eleven for longevity, thirteen for dispersal propensity and ten for sex ratio (Fig. 3B–Fig. 3G). Figure 3(B)–Figure 3(G) shows clear indications of a positive correlation between lifetime fecundity and its sixteen selected metabolites. In contrast, Figure 3(B)–Figure 3(G) suggests a negative correlation between the twenty and thirteen metabolites selected for, respectively, egg survival and dispersal propensity. For daily fecundity, longevity and sex ratio, no clear trends were visible.

In the subsequent linear regressions, one significant correlation was found for egg survival (a sugar alcohol) and seven for dispersal propensity (including four essential amino acids and one sugar). No significant results were found for lifetime fecundity, daily fecundity, longevity and sex ratio (Fig. 3B–Fig. 3G and appendix A.6).

Pathway analysis indicated that four of the seven metabolites which negatively correlated with dispersal propensity play a significant role in the aminoacyl-tRNA biosynthesis (total: 67, hits: 4, impact=0, Holm p=0.0417) (the pathway map is provided in appendix A.7). No associated pathways were found for egg survival.

## Discussion

Of the forty-three metabolites identified in the GC-MS analysis, eighteen correlated with latitude and/or one or more life-history traits. More specifically, eleven covaried positively or negatively with latitude, seven showed a negative correlation with dispersal propensity and one showed a negative correlation with egg survival. Of the eighteen different metabolites, eleven amino acids could be shown to play an important role in the aminoacyl-tRNA biosynthesis and four in the valine, leucine and isoleucine biosynthesis (see pathway maps provided in appendix A.7).

Contrary to our hypothesis, our results indicate that the life-history evolution which occurred during the recent range expansion of *T. urticae* (Van Petegem *et al.* 2016) was not associated with shifts in the mites’ energetic metabolism, but rather with shifts in its anabolism. While our spectral database contained eleven sugars, only one sugar (fructose) accounted for the separation among populations. This suggests that the genes involved in encoding the mite’s energetic metabolism (*i.e.* glycolysis, citric acid cycle, which typically involve sugars) have not been significantly affected during the range expansion of *T. urticae*. Instead, the observed differentiation in the mites’ metabolome probably involved evolutionary changes in the mites’ anabolism, where amino acids play a central role in the metabolic turnover of proteins. In more northern and more dispersive populations, the aminoacyl-tRNA biosynthesis was downregulated. In this pathway, aminoacyl-tRNA is formed by charging tRNA with an amino acid. The aminoacyl-tRNA then serves as a substrate in protein synthesis or plays one of its many other roles in, for example, cell wall formation or antibiotic biosynthesis (Raina & Ibba 2014). In accordance, the valine, leucine and isoleucine biosynthesis, important for protein synthesis as well (Ahmed & Khan 2006; Tamanna & Mahmood 2014), was downregulated in more northern populations.

The affected amino acids showed decreased concentrations toward higher latitudes and showed a negative correlation with the dispersal propensity of *T. urticae*, which increases towards the north. While, in general, amino acids are considered fundamental for egg production and thus fecundity (Tulisalo 1971; O'Brien, Fogel & Boggs 2002; Mevi-Schutz & Erhardt 2005; Fuchs *et al.* 2014, but see Heagle *et al.* 2002), not a single correlation was found for fecundity, despite a clear positive trend in the PLS-DA. Of the eleven affected amino acids, eight were essential and three non-essential. While the non-essential amino acids could have been synthesised *de novo* from glucose, the essential amino acids could only have been supplied through the mite's diet (Rodriguez *et al.* 1966). Though all mites were kept in common garden, mites from northern, more dispersive populations were found to contain lower essential amino acid concentrations. In line with the recent finding of Fronhofer & Altermatt (2015) that a dispersal-foraging trade-off leads to a reduced exploitation of resources at range margins, our results could indicate that northern, more dispersive mites evolved lower essential amino acid concentrations because they consume less of their food source. We should, however, keep in mind that metabolites were measured only at one point in time and from whole organism samples. We are therefore missing the temporal fluctuations of the metabolome over a day, and our data therefore represent only a snapshot of the existing balance in terms of metabolite demand among metabolic pathways.

An important challenge for metabolomics is understanding the relative contribution of environmental and genetic factors in shaping an organism’s metabolic phenotype (Bundy *et al.* 2009). In the current study, mites were kept in common garden for two generations, during which they were reared under optimal, non-stressful conditions. In contrast with most metabolomic studies, mites were thus not subjected to a stressor to elicit a strong environmentally-induced plastic stress-response. Therefore, any metabolomic differentiation was expected to be far less pronounced compared to stress-exposure studies. As plastic, environmentally driven field-differences among populations were largely levelled out through the two generations in common garden, only genetic factors were retained. Long-lasting transgenerational plasticity can, however, not be fully excluded and we therefore refer further to (epi)genetic factors (Verhoeven et al. 2016). Though (epi)genetic factors are generally considered less determining than environmental factors (Robinson *et al.* 2007; Frank, Noerenberg & Engel 2009; Matsuda *et al.* 2012), our results demonstrate a clear (epi)genetic signal of metabolic differentiation along *T. urticae’s* invasion gradient. We acknowledge, however, that we cannot exclude neutral processes, like for example serial bottlenecks-including surfing mutations (Travis *et al.* 2007; Klopfstein *et al.* 2016) as potential (co)sources of the found latitudinal metabolomic patterns. Furthermore, as the host plant species on which the mites were sampled in the field covaried with latitude (with *L. periclymenum* typically in the north), this could also have influenced our results. Yet, regression slopes barely differed between models including all populations (hence all host plant species) or models run for the subset of six populations collected on *L. periclymenum* only (see appendix A.8). This indicates that the latitudinal signal was independent of host plant identity.

This explorative study specifically examined whether range expansion might result in evolutionary changes in an organism’s metabolism. Despite non-stressful common garden conditions, approximately forty per cent of the identified metabolites showed (epi)genetic differentiation among populations. The more dispersive northern mites exhibited lower concentrations of several essential and non-essential amino acids, suggesting a downregulation of certain pathways linked to protein synthesis. Though effects were subtle (but see earlier), our results clearly indicate that the metabolome of *T. urticae* underwent (epi)genetic changes during the species’ recent range expansion.

## Acknowledgements

This project was funded by the Fund for Scientific Research-Flanders (FWO) (project G.0610.11). DB and RS were supported by BelSpo IAP Project “Speedy”. We furthermore thank the INEE-CNRS (ENVIROMICS call, project ‘ALIENS’) for funding DR.

## Data accessibility

– Sample locations and life-history trait values: included in appendix
– Results of GC-MS analysis: will be deposited in the Dryad repository (and added to the reference list)
– Outcomes of statistical analyses: included in appendix

## Appendix A.l: Overview of the Field Collection Sites and their Respective Life-History Trait Values

An overview of the localities where *T. urticae* was sampled in the field, together with population-level life-history trait values (daily fecundity (#eggs/day), lifetime fecundity (total #eggs), egg survival (%), longevity (#days), dispersal propensity (%) and sex ratio (#male/total)). These values, originating from a previous study (Van Petegem *et al.* 2016), were used as the independent values in our linear regressions. Field collection sites were located along a latitudinal gradient, spanning the coasts of Belgium (BEL), The Netherlands (NTL), Germany and Denmark (DEN). The denoted plant species (*Lonicera periclymenum, Euonymus europaeus, Sambucus nigra* and *Humulus lupulus*) are the species on which the mites were found and sampled.

| code | collection site | city, country | Coordinates<br>(WGS 84) | plant species | daily<br>fecundity | lifetime<br>fecundity | egg<br>survival | longevity | dispersal<br>propensity | sex<br>ratio |
| --- | --- | --- | --- | --- | --- | --- | --- | --- | --- | --- |
| SKA | Flagbakkevej | Skagen, DEN | 57.72, 10.53 | *L. periclymenum* | 3.85 | 33.42 | 96.72 | 10.80 | 9.47 | 0.33 |
| SVI | Sletteåvej | Fjerritslev, DEN | 57.15, 9.33 | *L. periclymenum* | 5.19 | 29.33 | . | 6.33 | 41.41 | 0.32 |
| TVE | Hindingvej | Thisted, DEN | 57.04, 8.62 | *L. periclymenum* | 3.65 | 23.42 | 100.00 | 7.42 | 14.53 | 0.26 |
| BLA | V. Hennebysvej | Henne, DEN | 55.74, 8.22 | *L. periclymenum* | 3.72 | 40.92 | 100.00 | 11.00 | 39.47 | 0.29 |
| HED | picnic-place along 11/24 | Gredstedbro, DEN | 55.39, 8.74 | *L. periclymenum* | 3.48 | 29.17 | 96.88 | 9.60 | 31.45 | 0.22 |
| LAU | Schildhoek | Lauwersoog, NTL | 53.41, 6.22 | *E. europaeus* | 4.25 | 37.00 | 87.30 | 9.75 | 16.75 | 0.34 |
| CAS | Hoofdweg | Heemskerk, NTL | 52.53, 4.65 | *L. periclymenum* | 5.54 | 41.75 | 96.15 | 9.40 | 19.21 | 0.28 |
| KVS | Bosweg | Burgh-Haamstede, NTL | 51.68, 3.72 | *S. nigra* | 6.34 | 98.67 | 99.02 | 17.55 | 15.03 | 0.19 |
| ODK | Duinparklaan | Koksijde, BEL | 51.12, 2.68 | *H. lupulus* | 5.22 | 50.42 | 98.99 | 10.08 | 9.81 | 0.25 |

## A.2: Overview of the 60 Metabolites Included in Our Spectral Database

A list of all 60 metabolites screened for in our single quadrupole GC-MS. Each of these metabolites can be characterised by several ions of different masses. We used the selective ion monitoring mode (SIM, see Waller *et al.* 2007) instead of the full scan mode. In this SIM mode, the MS instrument is set to only look for specific previously-known ion masses of interest rather than to screen for all masses over a wide range. The instrument can therefore be very specific for a particular compound of interest. Our original database contained spectral information for 85 primary metabolites from a range of plant and invertebrate models. After removing all metabolites specific to plant models and those (polyamines for instance) having concentrations lower than the detection limit of our equipment, only the 60 specific metabolites listed in the table below remained in our spectral database. Metabolites are classified according to six categories: amino acid, polyol, sugar, intermediate of the citric acid cycle,‘other’ metabolite and metabolite not found in any of our samples.

| category | metabolite |
| --- | --- |
| amino acid | Alanine |
| amino acid | Aspartic acid |
| amino acid | Glutamic acid |
| amino acid | Glycine |
| amino acid | Isoleucine |
| amino acid | Leucine |
| amino acid | Lysine |
| amino acid | Methionine |
| amino acid | Phenylalanine |
| amino acid | Proline |
| amino acid | Serine |
| amino acid | Threonine |
| amino acid | Tryptophan |
| amino acid | Tyrosine |
| amino acid | Valine |
| intermediate citric acid cycle | Citric acid |
| intermediate citric acid cycle | Fumaric acid |
| intermediate citric acid cycle | Malic acid |
| intermediate citric acid cycle | Succinic acid |
| polyol | Adonitol |
| polyol | Arabitol |
| polyol | Glycerol |
| polyol | Glycerol-3-phosphate |
| polyol | Inositol |
| polyol | Sorbitol |
| polyol | Xylitol |
| sugar | Fructose |
| sugar | Fructose-6-phosphate |
| sugar | Galactose |
| sugar | Glucose |
| sugar | Glucose-6-phosphate |
| sugar | Mannose |
| sugar | Ribose |
| other | Ethanolamine |
| other | GABA |
| other | Gluconolactone |
| other | Glyceric acid |
| other | Lactic acid |
| other | Ornithine |
| other | Phosphoric acid |
| other | Putrescine |
| other | Quinic acid |
| other | Spermidine |
| not found | Arabinose |
| not found | Asparagine |
| not found | Cadaverine |
| not found | Citrulline |
| not found | Cytosine |
| not found | Erythritol |
| not found | Galactitol |
| not found | Galacturonic acid |
| not found | Maltose |
| not found | Mannitol |
| not found | Octopamine |
| not found | Pipecolic acid |
| not found | Saccharose |
| not found | Spermine |
| not found | Trehalose |
| not found | Tyramine |
| not found | Xylose |

## A.3: Correlation Matrix

LAT (latitude), LIFE (lifetime fecundity), DAFE (daily fecundity), DISP (dispersal propensity), EGSU (egg survival), LONG (longevity), DIAP (diapause incidence) and SERA (sex ratio). All significant correlations (Bonferroni correction not implemented) are in bold. As only a selection of populations was included in the current study, the strength of the correlations between latitude and the shown life-history traits decreased relative to the correlations found in Van Petegem *et al.* (2016).

| Pearson Correlation Coefficients |  |  |  |  |  |  |  |  |
| --- | --- | --- | --- | --- | --- | --- | --- | --- |
| Prob > \|r\| under H0: Rho=0 |  |  |  |  |  |  |  |  |
| Number of Observations |  |  |  |  |  |  |  |  |
|  | LAT | LIFE | DAFE | DISP | EGSU | LONG | DIAP | SERA |
| LAT | 1 | -0.67384 | -0.67452 | 0.38101 | 0.13882 | -0.52688 | 0.47464 | 0.43955 |
|  |  | **0.0466** | **0.0463** | 0.3117 | 0.743 | 0.145 | 0.1967 | 0.2365 |
|  | 9 | 9 | 9 | 9 | 8 | 9 | 9 | 9 |
| LIFE | -0.67384 | 1 | 0.75065 | -0.27469 | 0.18641 | 0.9203 | -0.337 | -0.5922 |
|  | **0.0466** |  | **0.0198** | 0.4744 | 0.6585 | **0.0004** | 0.3752 | 0.0929 |
|  | 9 | 9 | 9 | 9 | 8 | 9 | 9 | 9 |
| DAFE | -0.67452 | 0.75065 | 1 | -0.18302 | 0.06919 | 0.46919 | 0.13277 | -0.33554 |
|  | **0.0463** | **0.0198** |  | 0.6374 | 0.8707 | 0.2026 | 0.7335 | 0.3774 |
|  | 9 | 9 | 9 | 9 | 8 | 9 | 9 | 9 |
| DISP | 0.38101 | -0.27469 | -0.18302 | 1 | 0.14537 | -0.31841 | 0.66023 | 0.11912 |
|  | 0.3117 | 0.4744 | 0.6374 |  | 0.7313 | 0.4037 | 0.0529 | 0.7602 |
|  | 9 | 9 | 9 | 9 | 8 | 9 | 9 | 9 |
| EGSU | 0.13882 | 0.18641 | 0.06919 | 0.14537 | 1 | 0.16269 | 0.19129 | -0.59346 |
|  | 0.743 | 0.6585 | 0.8707 | 0.7313 |  | 0.7003 | 0.65 | 0.1209 |
|  | 8 | 8 | 8 | 8 | 8 | 8 | 8 | 8 |
| LONG | -0.52688 | 0.9203 | 0.46919 | -0.31841 | 0.16269 | 1 | -0.52876 | -0.54739 |
|  | 0.145 | **0.0004** | 0.2026 | 0.4037 | 0.7003 |  | 0.1433 | 0.1271 |
|  | 9 | 9 | 9 | 9 | 8 | 9 | 9 | 9 |
| DIAP | 0.47464 | -0.337 | 0.13277 | 0.66023 | 0.19129 | -0.52876 | 1 | 0.29697 |
|  | 0.1967 | 0.3752 | 0.7335 | 0.0529 | 0.65 | 0.1433 |  | 0.4377 |
|  | 9 | 9 | 9 | 9 | 8 | 9 | 9 | 9 |
| SERA | 0.43955 | -0.5922 | -0.33554 | 0.11912 | -0.59346 | -0.54739 | 0.29697 | 1 |
|  | 0.2365 | 0.0929 | 0.3774 | 0.7602 | 0.1209 | 0.1271 | 0.4377 |  |
|  | 9 | 9 | 9 | 9 | 8 | 9 | 9 | 9 |

## A.4: 3-D Score Plots

3-D score plots resulting from the multivariate analyses (PLS-DA) performed on our metabolomic data are given for latitude (A) and each of six life-history traits (daily fecundity (B), lifetime fecundity (C), egg survival (D), longevity (E), dispersal propensity (F) and sex ratio (G)). The mean metabolic phenotypes for each replicate (five in total) of all nine populations (eight for egg survival) are represented by a dot and arranged in space according to their projections on three component axes. Each of these axes explains a certain amount (%) of the metabolic variation that is present in the dataset. More similar points (populations with a similar metabolome) are placed closer together.

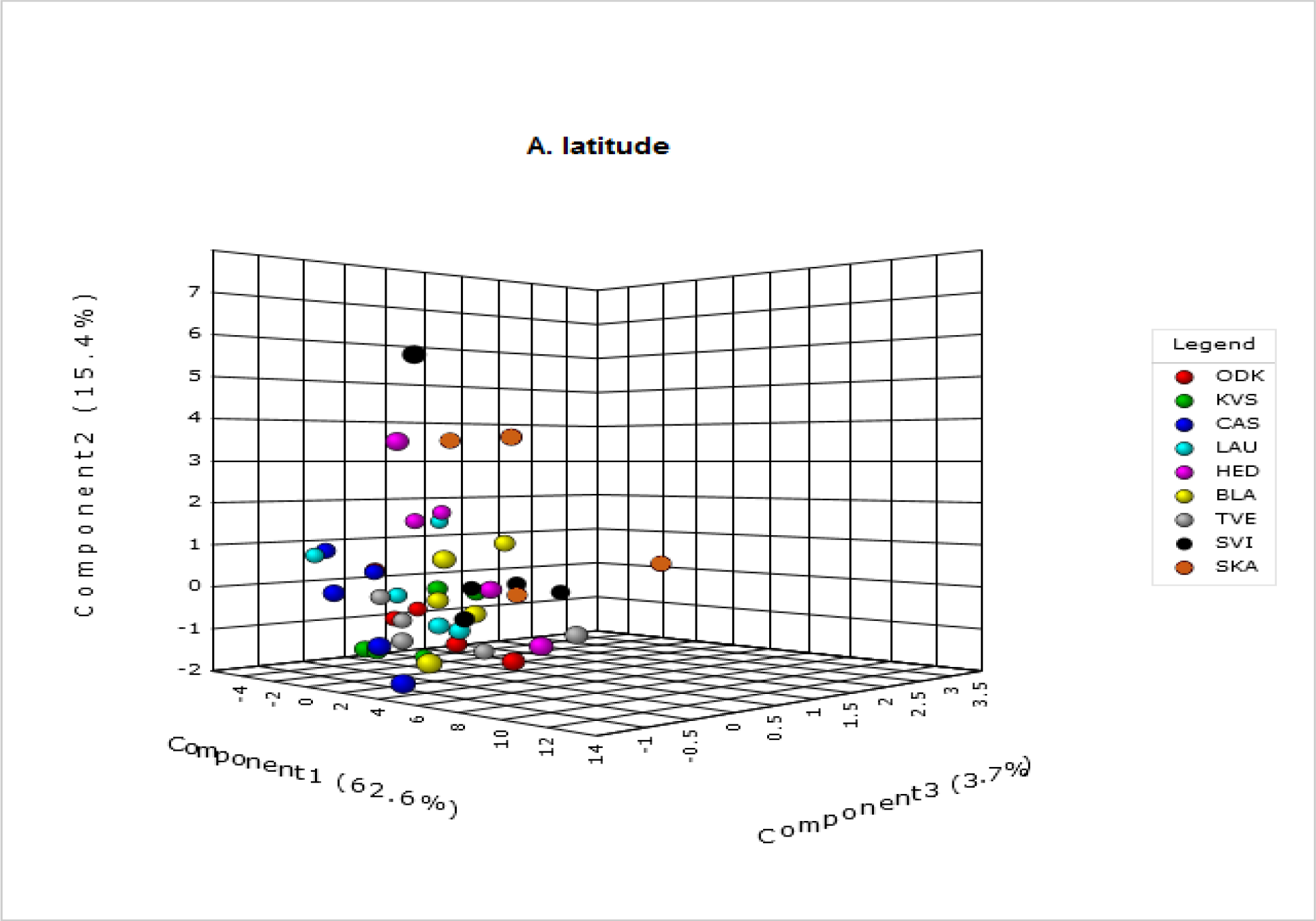

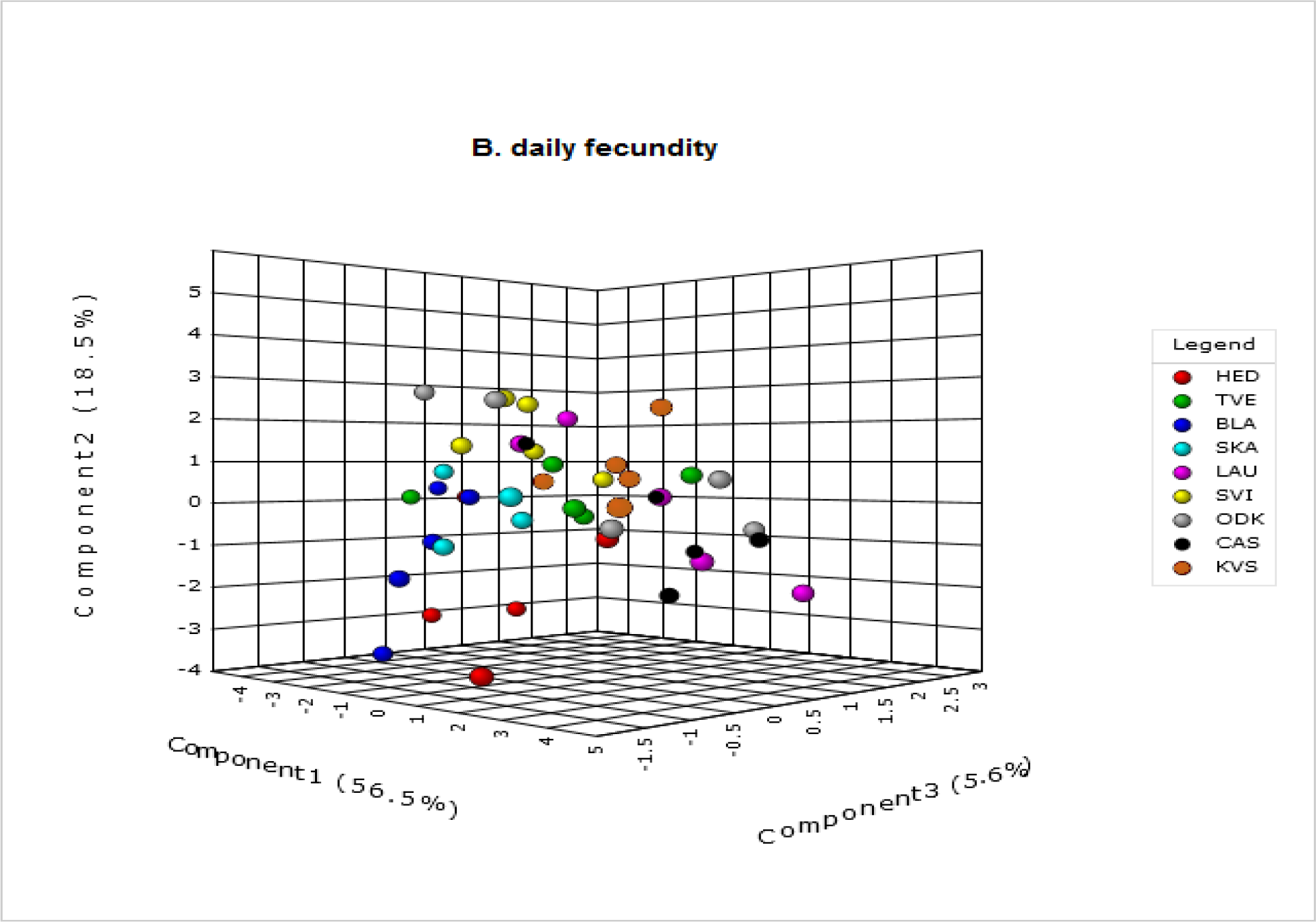

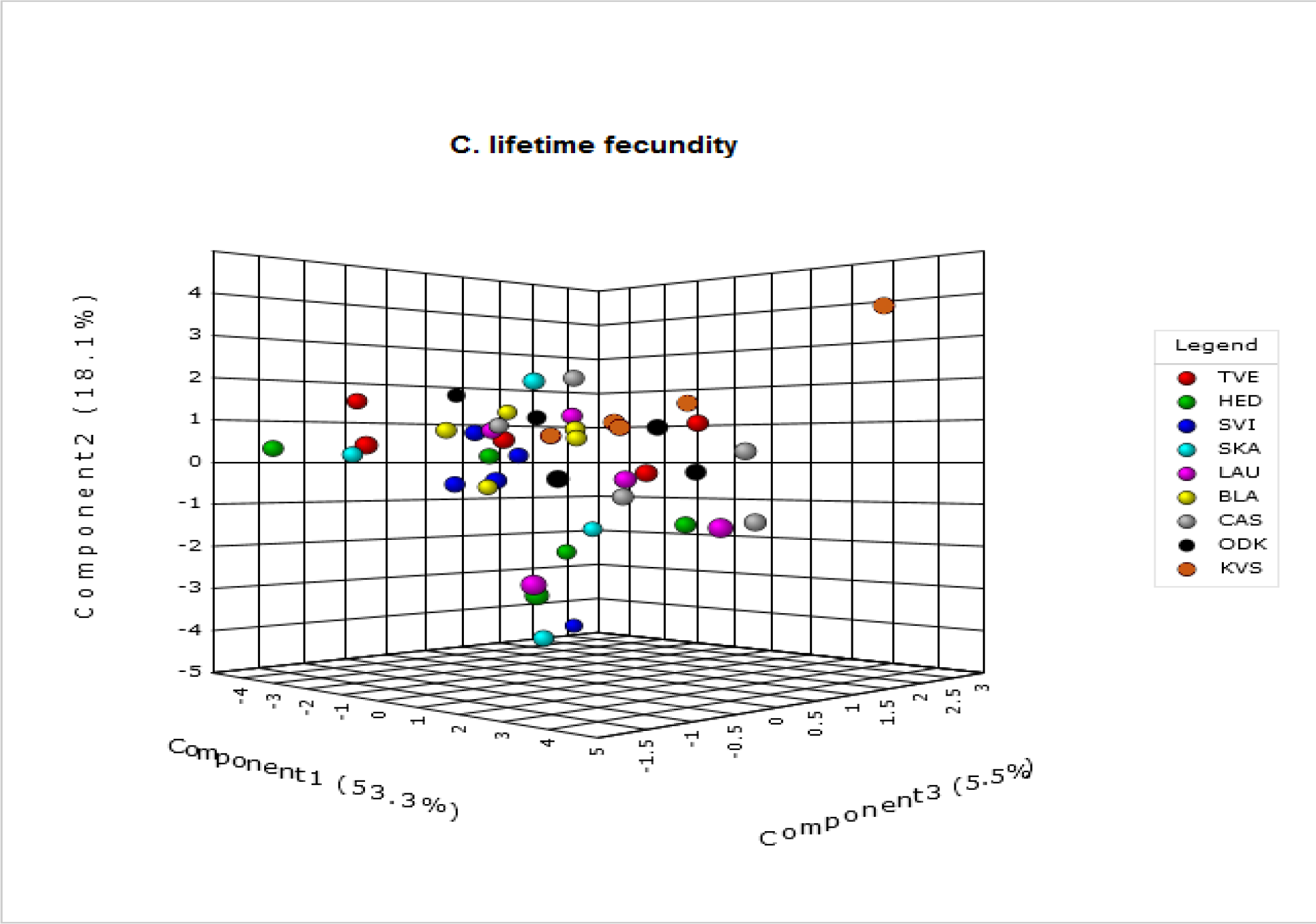

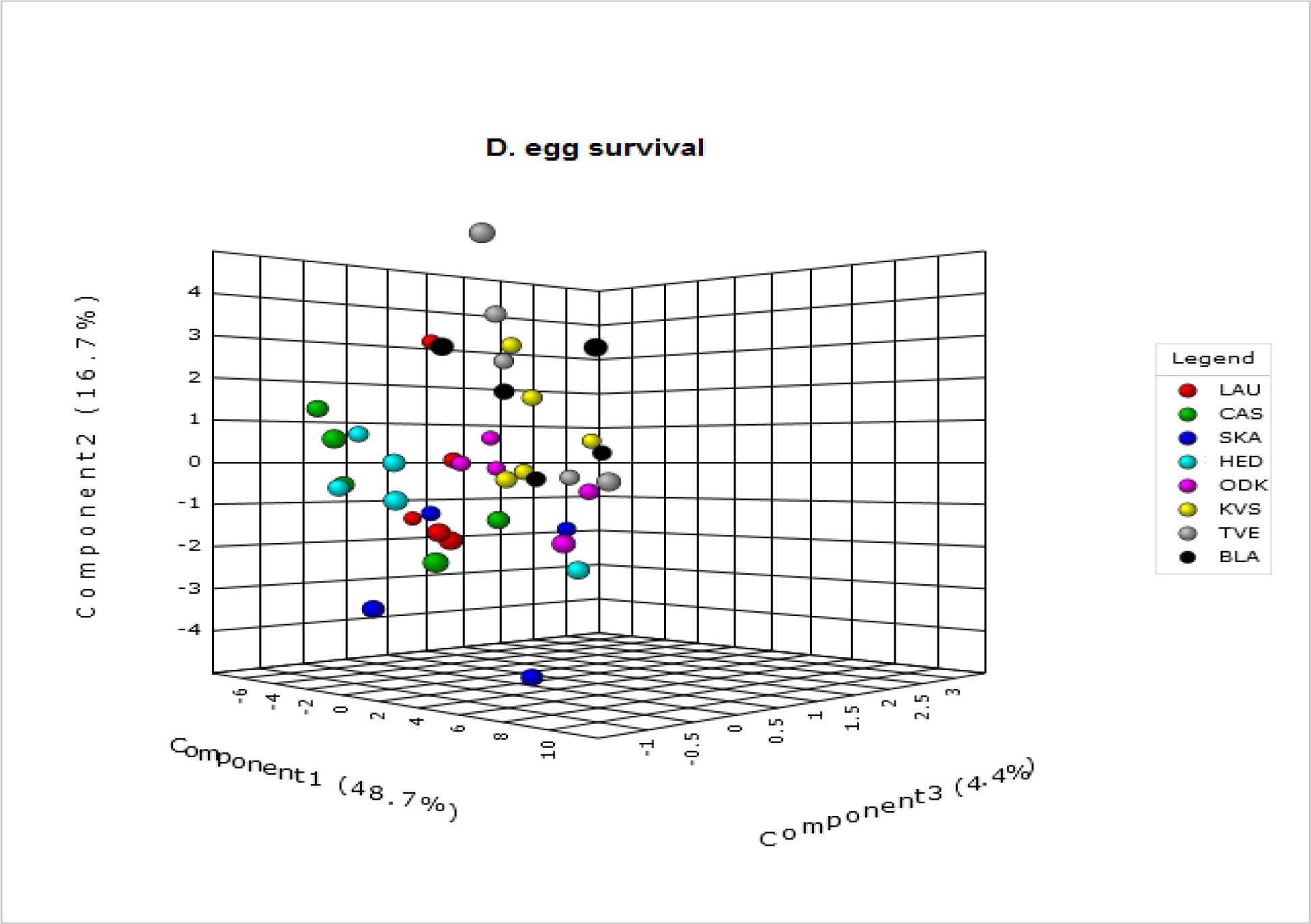

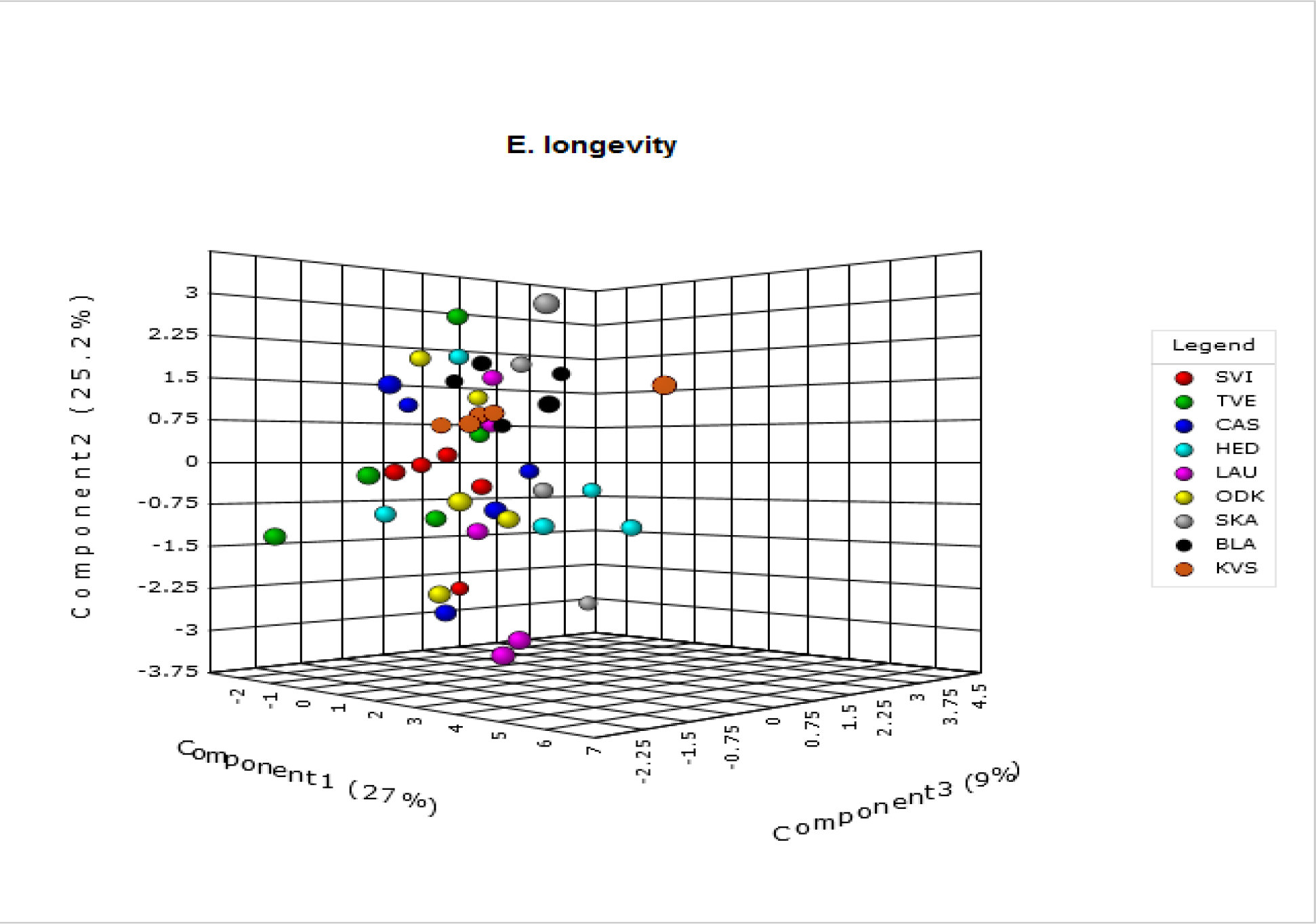

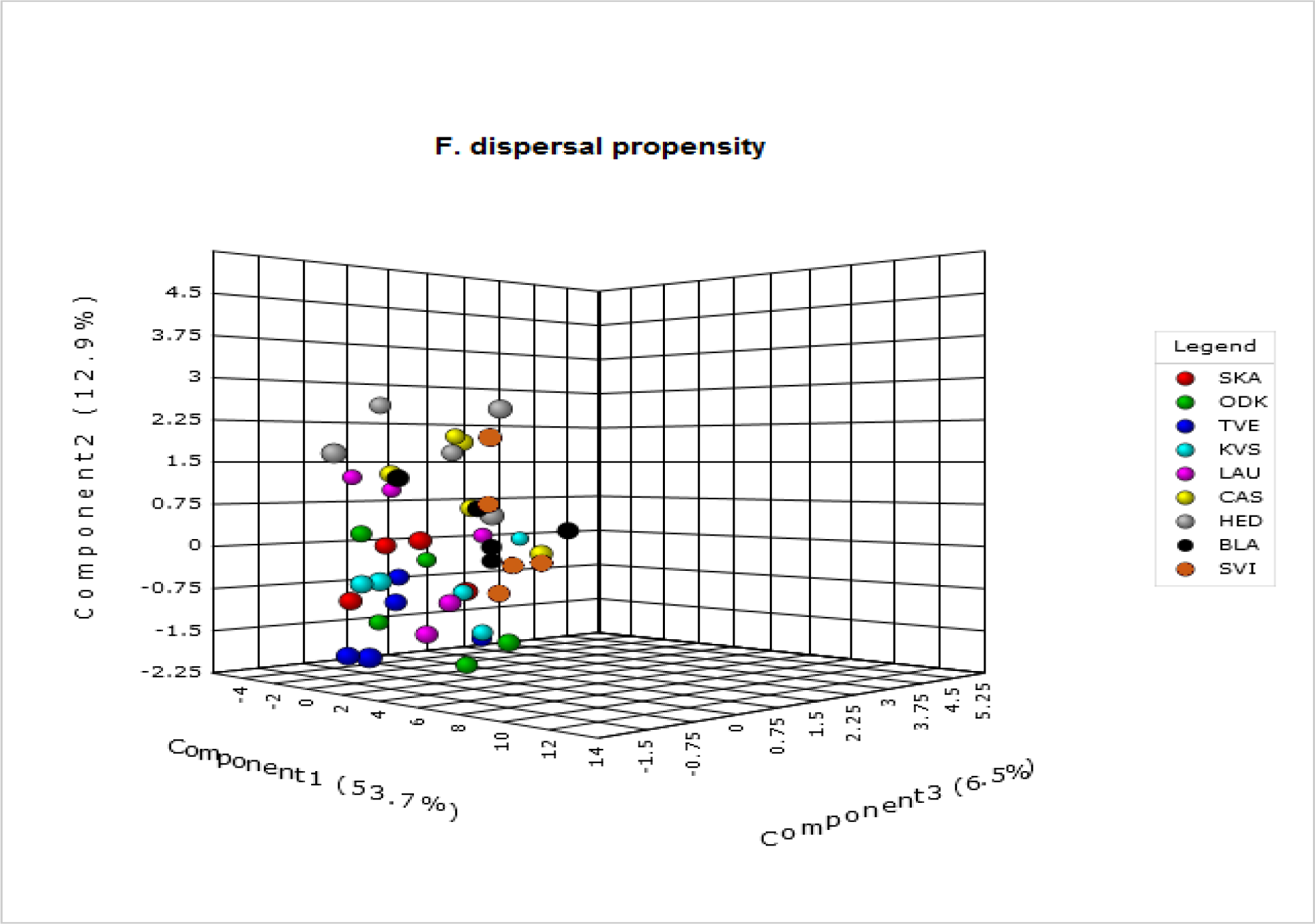

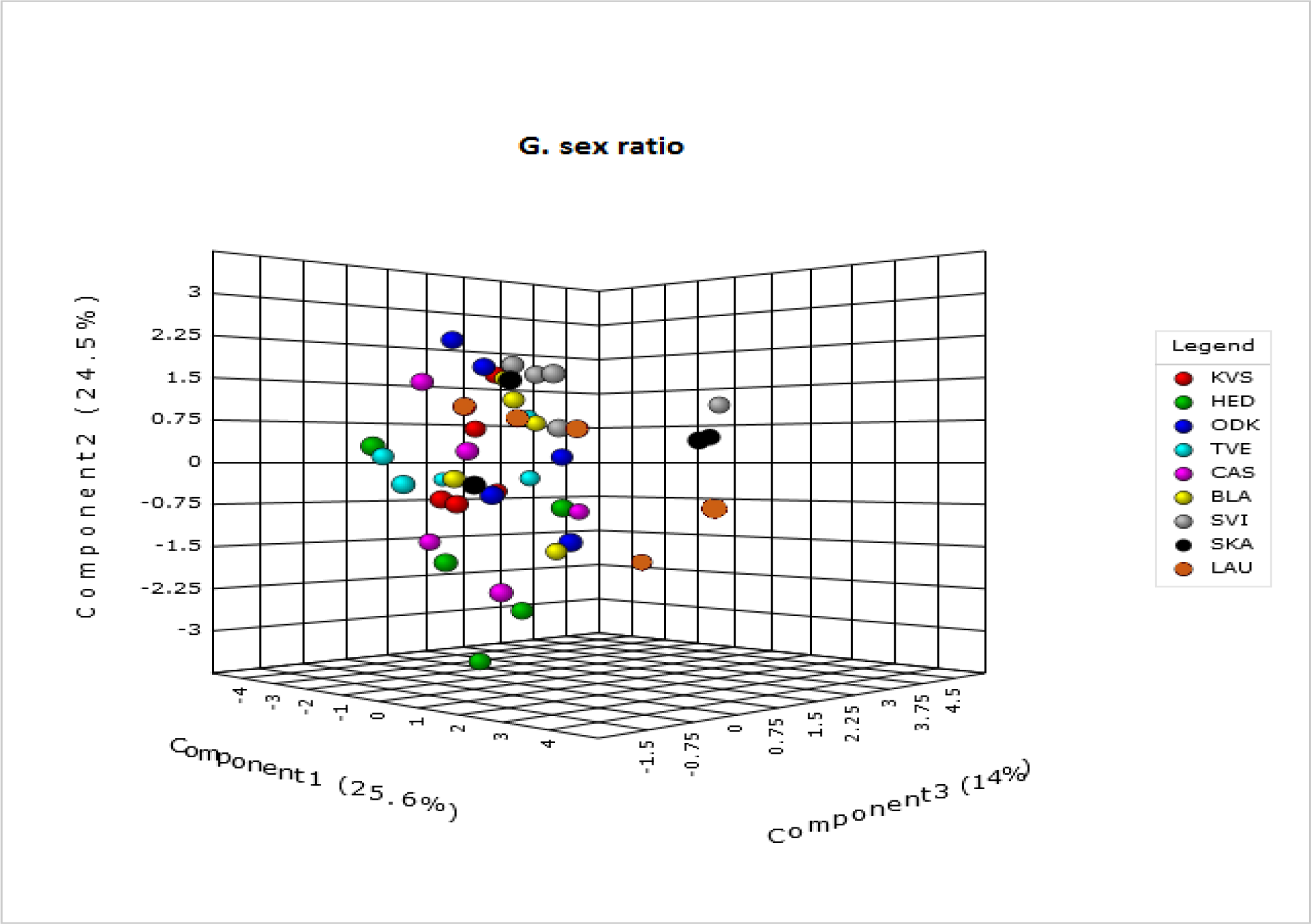

## A.5 Overview of the VIP Scores Resulting from the PLS-DA

An overview off all metabolites with a VIP score of at least 1.2 (1.0 for egg survival because removing the metabolites with a score between 1.0 and 1.2 resulted in a decreased percentage of variation explained) for component 1 and/ or 2 in the PLS-DA performed for latitude (A) and each of six life-history traits (daily fecundity (B), lifetime fecundity (C), egg survival (D), longevity (E), dispersal propensity (F) and sex ratio(G)). Note that in the tables below, the VIP scores for the original dataset (containing all forty-three identified metabolites) are given. The order of the metabolites (from high to low VIP scores) in these tables can therefore deviate from the order in Fig. 3 (main text), as Fig. 3 is based on VIP scores of the final dataset (after removal of all non-explanatory metabolites).

**A.** latitude

A. latitude
| metabolite | Comp. 1 | Comp. 2 |
| --- | --- | --- |
| Proline | 1.6583 | 1.0501 |
| Tryptophan | 1.5937 | 1.0289 |
| Inositol | 1.5831 | 1.0274 |
| Malic acid | 1.4947 | 0.9356 |
| Lysine | 1.4669 | 0.9290 |
| Valine | 1.4577 | 0.9261 |
| Isoleucine | 1.4219 | 0.9054 |
| Methionine | 1.4115 | 0.8879 |
| Glycine | 1.4090 | 0.8936 |
| Aspartic acid | 1.3881 | 0.8733 |
| Threonine | 1.3705 | 0.9118 |
| Alanine | 1.3572 | 0.8500 |
| Leucine | 1.3414 | 0.9084 |
| Quinic acid | 1.3313 | 1.6940 |
| Glucose-6-phosphate | 0.6854 | 2.0669 |
| Sorbitol | 1.0691 | 1.9186 |
| Fructose-6-phosphate | 0.4123 | 1.7967 |
| Ribose | 0.8471 | 1.2669 |

**B.**
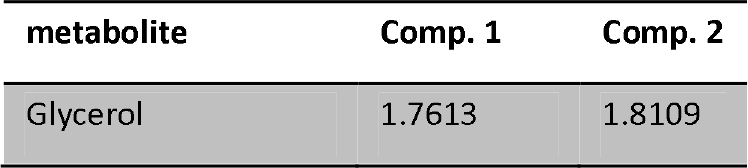
daily fecundity

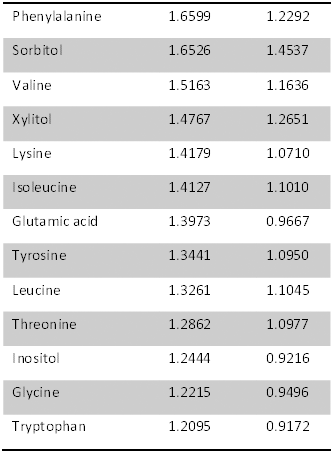

**C.**
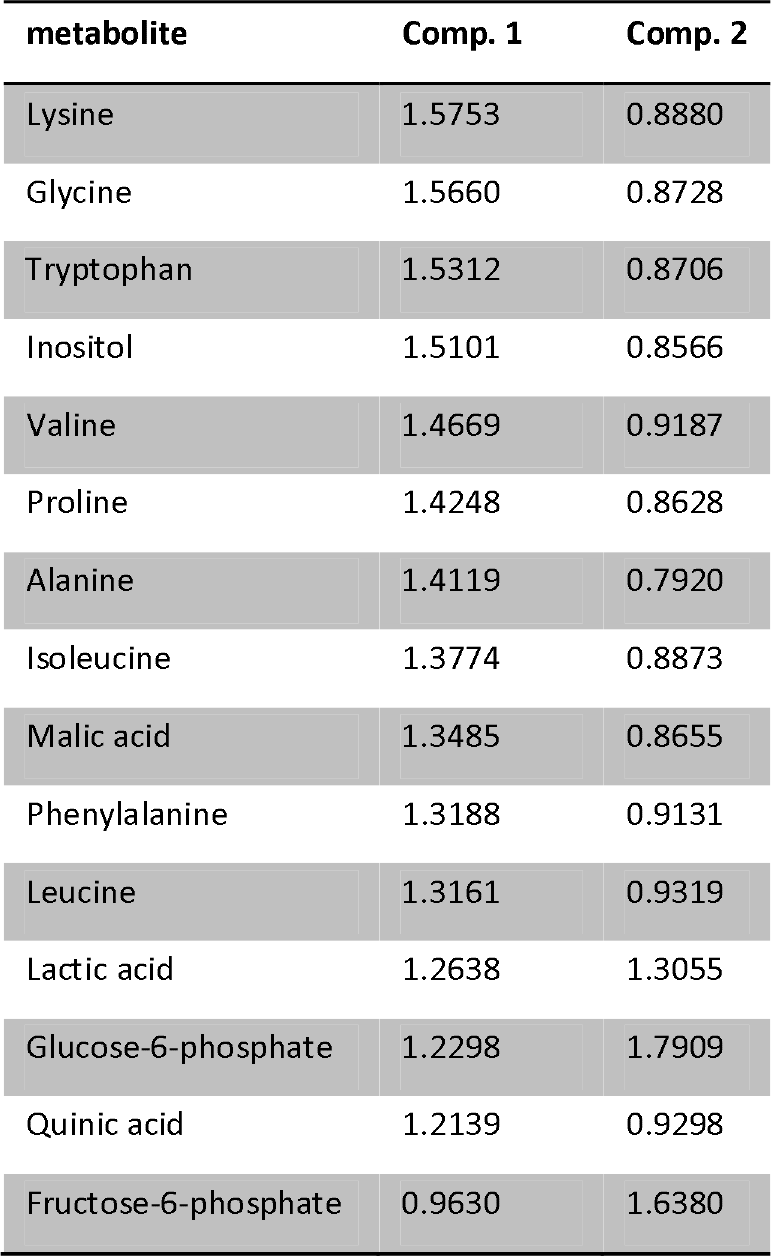
lifetime fecundity

C. lifetime fecundity
| metabolite | Comp. 1 | Comp. 2 |
| --- | --- | --- |
| Lysine | 1.5753 | 0.8880 |
| Glycine | 1.5660 | 0.8728 |
| Tryptophan | 1.5312 | 0.8706 |
| Inositol | 1.5101 | 0.8566 |
| Valine | 1.4669 | 0.9187 |
| Proline | 1.4248 | 0.8628 |
| Alanine | 1.4119 | 0.7920 |
| Isoleucine | 1.3774 | 0.8873 |
| Malic acid | 1.3485 | 0.8655 |
| Phenylalanine | 1.3188 | 0.9131 |
| Leucine | 1.3161 | 0.9319 |
| Lactic acid | 1.2638 | 1.3055 |
| Glucose-6-phosphate | 1.2298 | 1.7909 |
| Quinic acid | 1.2139 | 0.9298 |
| Fructose-6-phosphate | 0.9630 | 1.6380 |

**D.** egg survival

D. egg survival
| metabolite | Comp. 1 | Comp. 2 |
| --- | --- | --- |
| Glutamic acid | 1.7874 | 1.3641 |
| Alanine | 1.6795 | 1.1028 |
| Tyrosine | 1.4029 | 0.9337 |
| Phenylalanine | 1.3828 | 0.9234 |
| Fructose-6-phosphate | 1.3581 | 0.8561 |
| Glucose-6-phosphate | 1.3492 | 0.8500 |
| Methionine | 1.2596 | 0.8576 |
| Xylitol | 1.2585 | 1.0527 |
| Glycerol-3-phosphate | 1.2193 | 0.7804 |
| Citric acid | 1.2164 | 0.8394 |
| Serine | 1.1977 | 0.9293 |
| GABA | 1.1951 | 2.2339 |
| Gluconolactone | 1.1857 | 0.8810 |
| Adonitol | 1.1823 | 0.8112 |
| Succinic acid | 1.1628 | 1.0936 |
| Leucine | 1.1035 | 0.9157 |
| Threonine | 1.0640 | 0.9255 |
| Valine | 1.0216 | 0.8728 |
| Ornithine | 1.0141 | 0.6404 |
| Ethanolamine | 0.8772 | 1.9297 |
| Glycerol | 0.5981 | 1.7154 |
| Fumaric acid | 0.0152 | 1.5159 |
| Glutamic acid | 1.7874 | 1.3641 |
| Glyceric acid | 2.1181 | 1.3224 |

**E.**
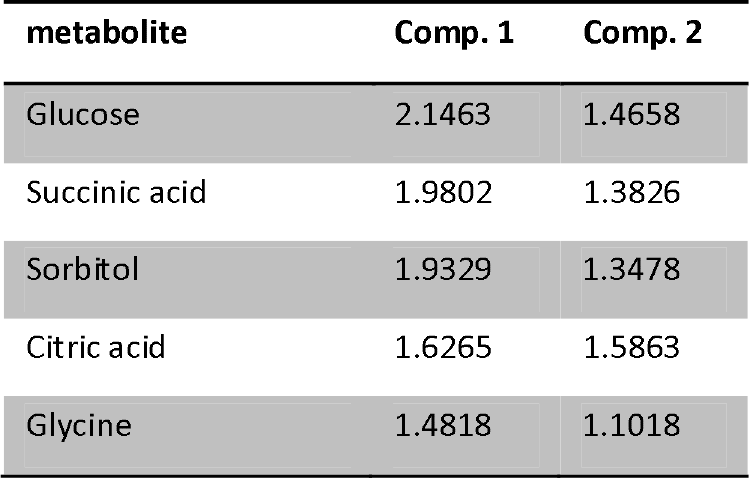
longevity

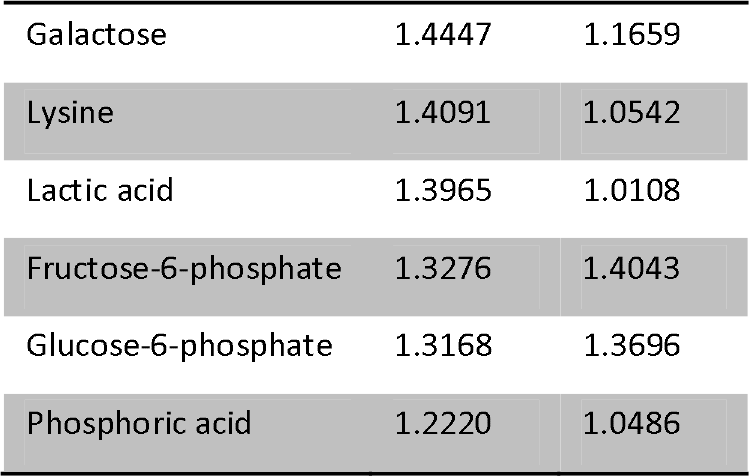

**F.** dispersal propensity

F. dispersal propensity
| metabolite | Comp. 1 | Comp. 2 |
| --- | --- | --- |
| Putrescine | 1.9187 | 1.2144 |
| Phosphoric acid | 1.7678 | 1.0416 |
| Lysine | 1.7503 | 1.0280 |
| Methionine | 1.7343 | 1.0303 |
| Xylitol | 1.7230 | 1.9394 |
| Phenylalanine | 1.5574 | 0.9821 |
| Fructose | 1.4964 | 0.9235 |
| Tryptophan | 1.2684 | 0.7950 |
| Glycerol-3-phosphate | 1.2602 | 0.7514 |
| Leucine | 1.2553 | 0.9363 |
| Inositol | 1.2549 | 0.7824 |
| Glucose-6-phosphate | 0.6820 | 1.4379 |
| Mannose | 0.2110 | 1.3947 |

**G.**
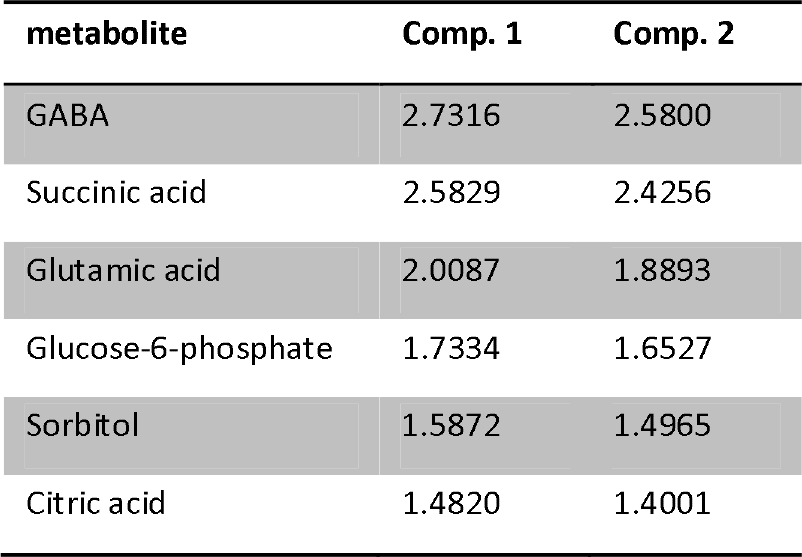
sex ratio

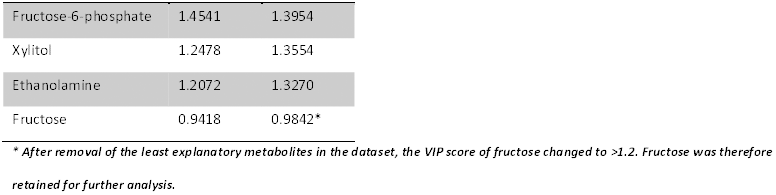

## A.6 Overview of all Linear Regressions

An overview of all linear regressions according to latitude (A) and each of the six life-history traits (daily fecundity (B), lifetime fecundity (C), egg survival (D), longevity (E), dispersal propensity (F) and sex ratio(G)) known to covary with latitude in the study species (see Van Petegem *et al.* 2016). The direction of change (correlation) is each time given, together with the F-and *p*-value of the linear regression. The degrees of freedom used in determining the F-value (Num DF and Den DF) are also shown.

**A.** latitude

A. latitude
| metabolite | correlation | Num DF | Den DF | F | p |
| --- | --- | --- | --- | --- | --- |
| Proline | negative | 1 | 7 | 9.85 | **0.0164** |
| Tryptophan | negative | 1 | 7 | 6.48 | **0.0383** |
| Inositol | negative | 1 | 7 | 6.43 | **0.0389** |
| Aspartic acid | negative | 1 | 7 | 6.62 | **0.0369** |
| Glycine | negative | 1 | 7 | 7.19 | **0.0315** |
| Malic acid | negative | 1 | 7 | 6.59 | **0.0372** |
| Valine | negative | 1 | 7 | 8.15 | **0.0245** |
| Lysine | negative | 1 | 7 | 3.77 | 0.0932 |
| Alanine | negative | 1 | 7 | 4.16 | 0.0808 |
| Methionine | negative | 1 | 7 | 3.06 | 0.1238 |
| Isoleucine | negative | 1 | 7 | 8.81 | **0.0209** |
| Quinic acid | positive | 1 | 7 | 5.59 | **0.0500** |
| Threonine | negative | 1 | 7 | 7.35 | **0.0301** |
| Leucine | negative | 1 | 7 | 6.71 | **0.0359** |
| Sorbitol | positive | 1 | 7 | 1.13 | 0.3221 |
| Glucose-6-phosphate | positive | 1 | 7 | 1.00 | 0.3511 |
| Fructose-6-phosphate | positive | 1 | 7 | 0.34 | 0.5789 |

**B.**
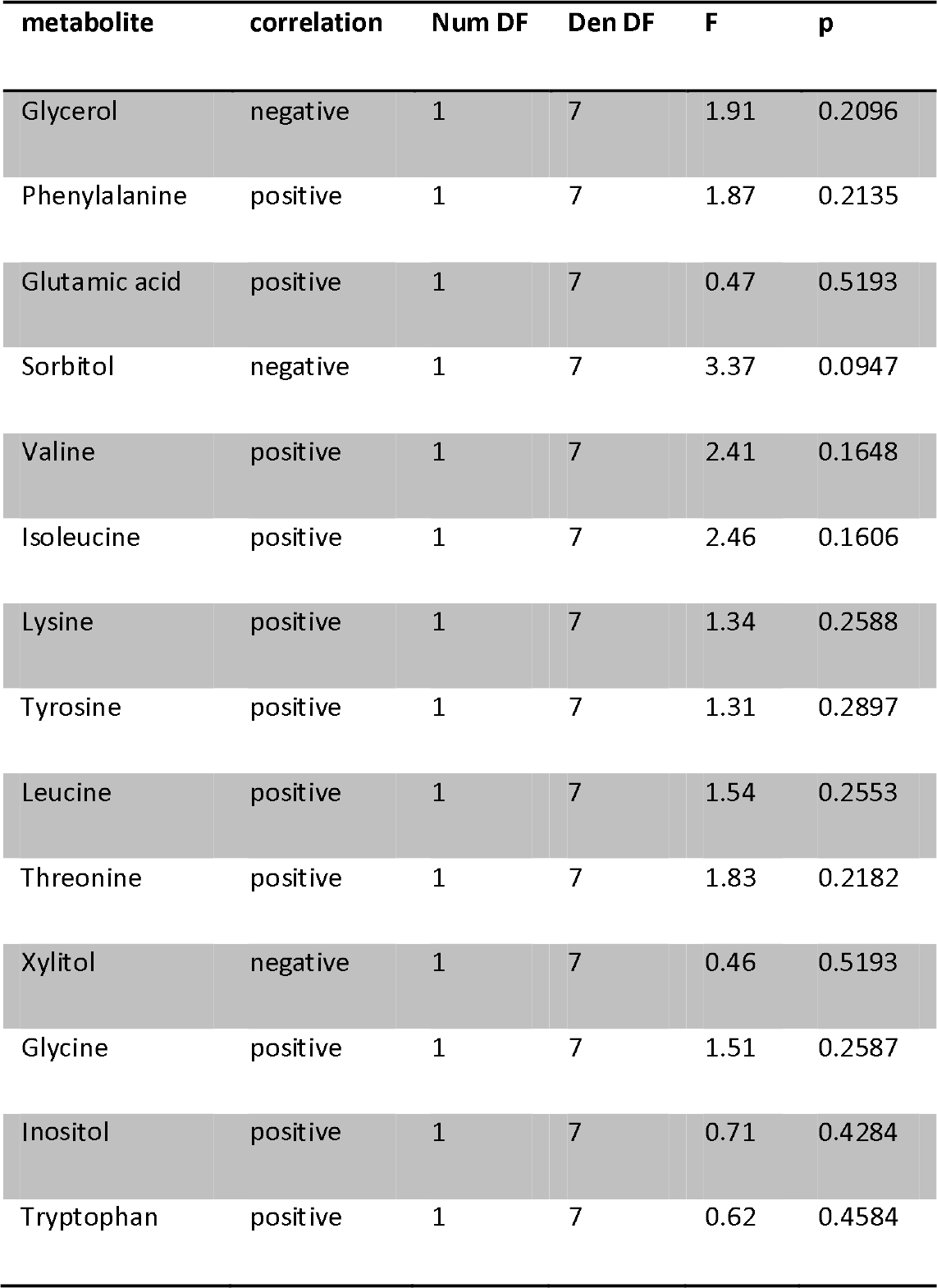
daily fecundity

B. daily fecundity
| metabolite | correlation | Num DF | Den DF | F | p |
| --- | --- | --- | --- | --- | --- |
| Glycerol | negative | 1 | 7 | 1.91 | 0.2096 |
| Phenylalanine | positive | 1 | 7 | 1.87 | 0.2135 |
| Glutamic acid | positive | 1 | 7 | 0.47 | 0.5193 |
| Sorbitol | negative | 1 | 7 | 3.37 | 0.0947 |
| Valine | positive | 1 | 7 | 2.41 | 0.1648 |
| Isoleucine | positive | 1 | 7 | 2.46 | 0.1606 |
| Lysine | positive | 1 | 7 | 1.34 | 0.2588 |
| Tyrosine | positive | 1 | 7 | 1.31 | 0.2897 |
| Leucine | positive | 1 | 7 | 1.54 | 0.2553 |
| Threonine | positive | 1 | 7 | 1.83 | 0.2182 |
| Xylitol | negative | 1 | 7 | 0.46 | 0.5193 |
| Glycine | positive | 1 | 7 | 1.51 | 0.2587 |
| Inositol | positive | 1 | 7 | 0.71 | 0.4284 |
| Tryptophan | positive | 1 | 7 | 0.62 | 0.4584 |

**C.**
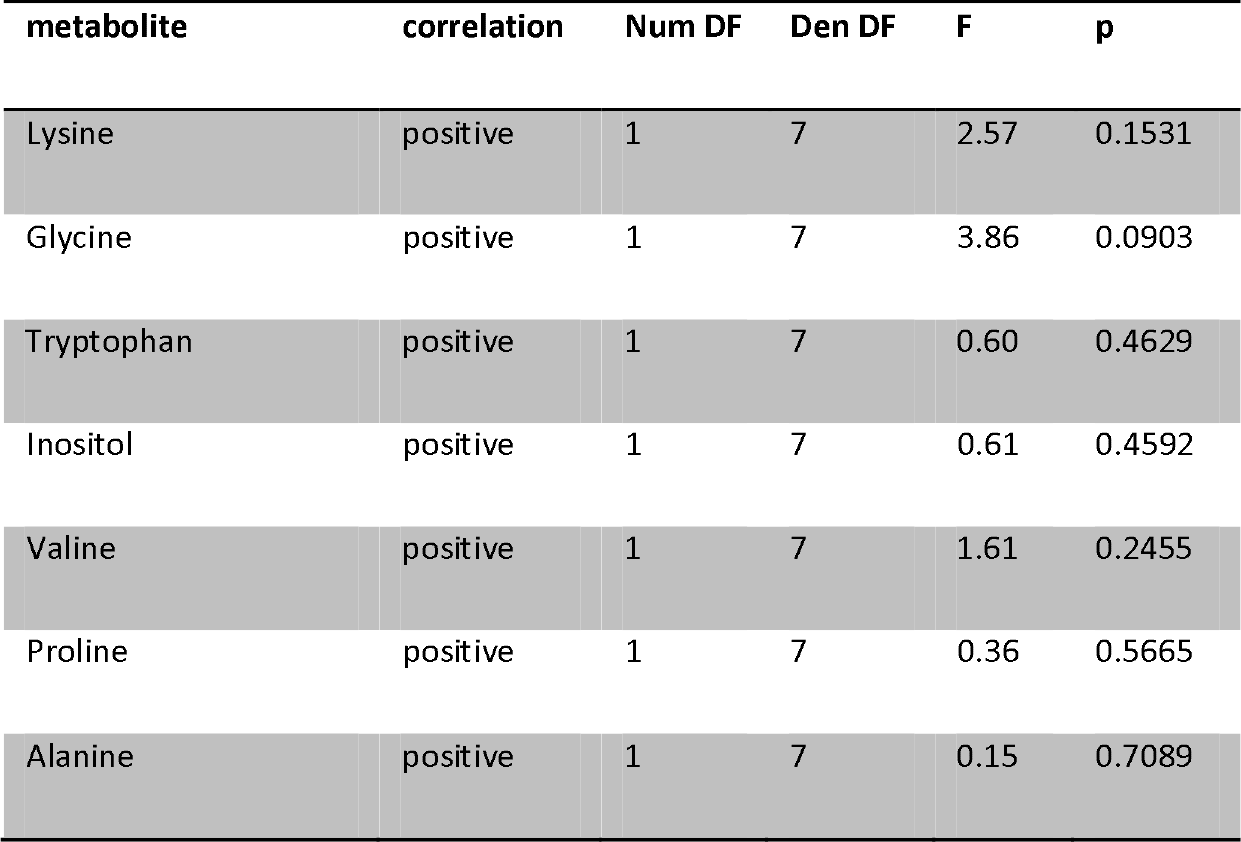
lifetime fecundity

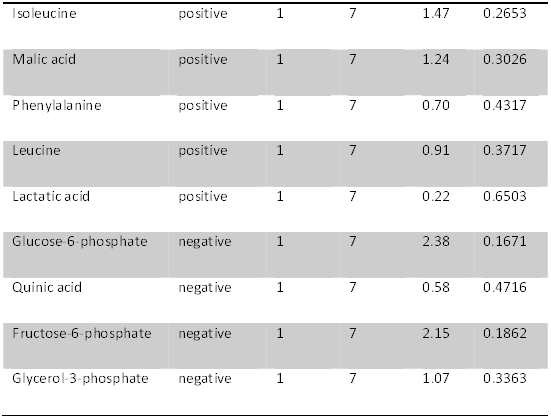

**D.**
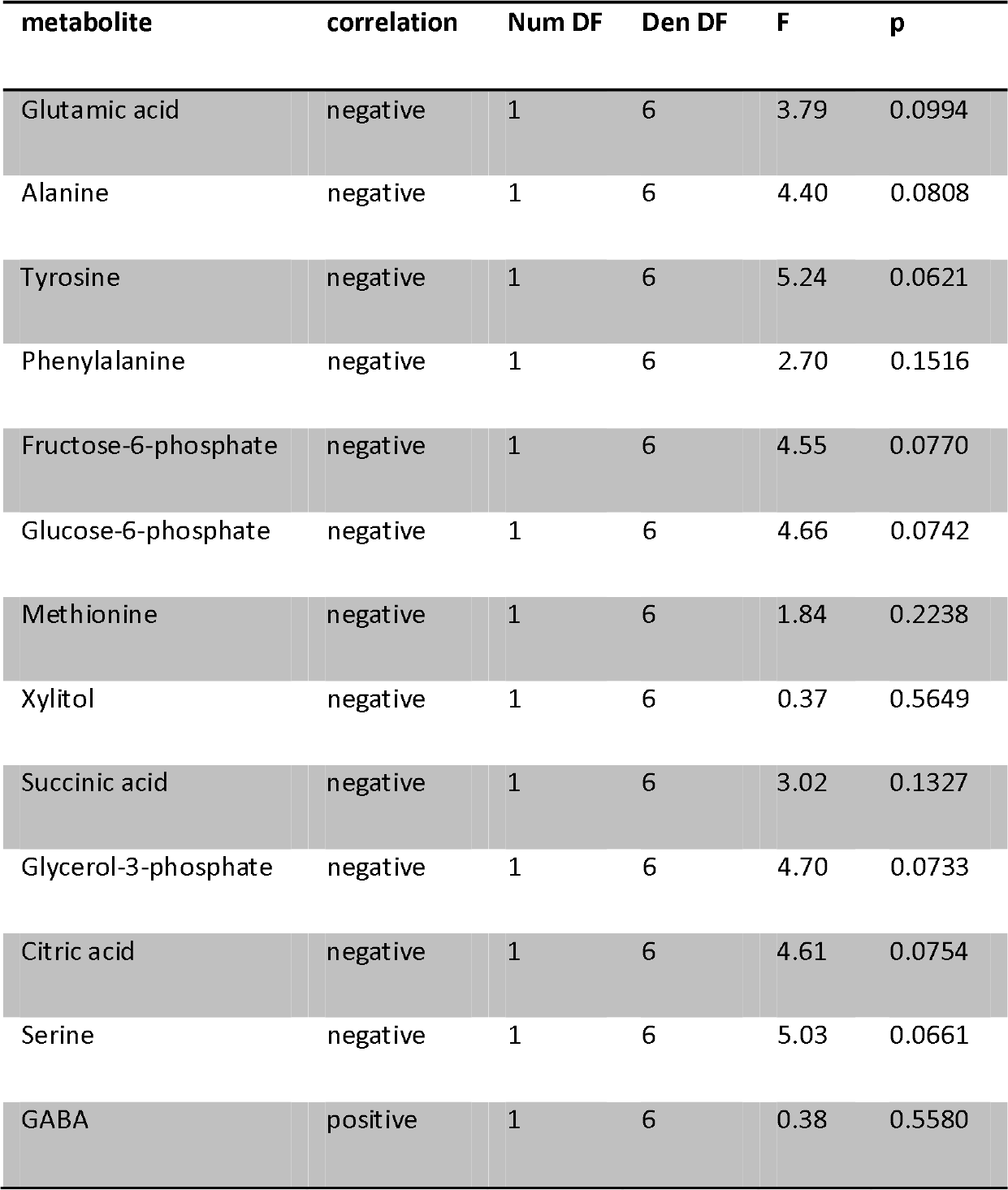
egg survival

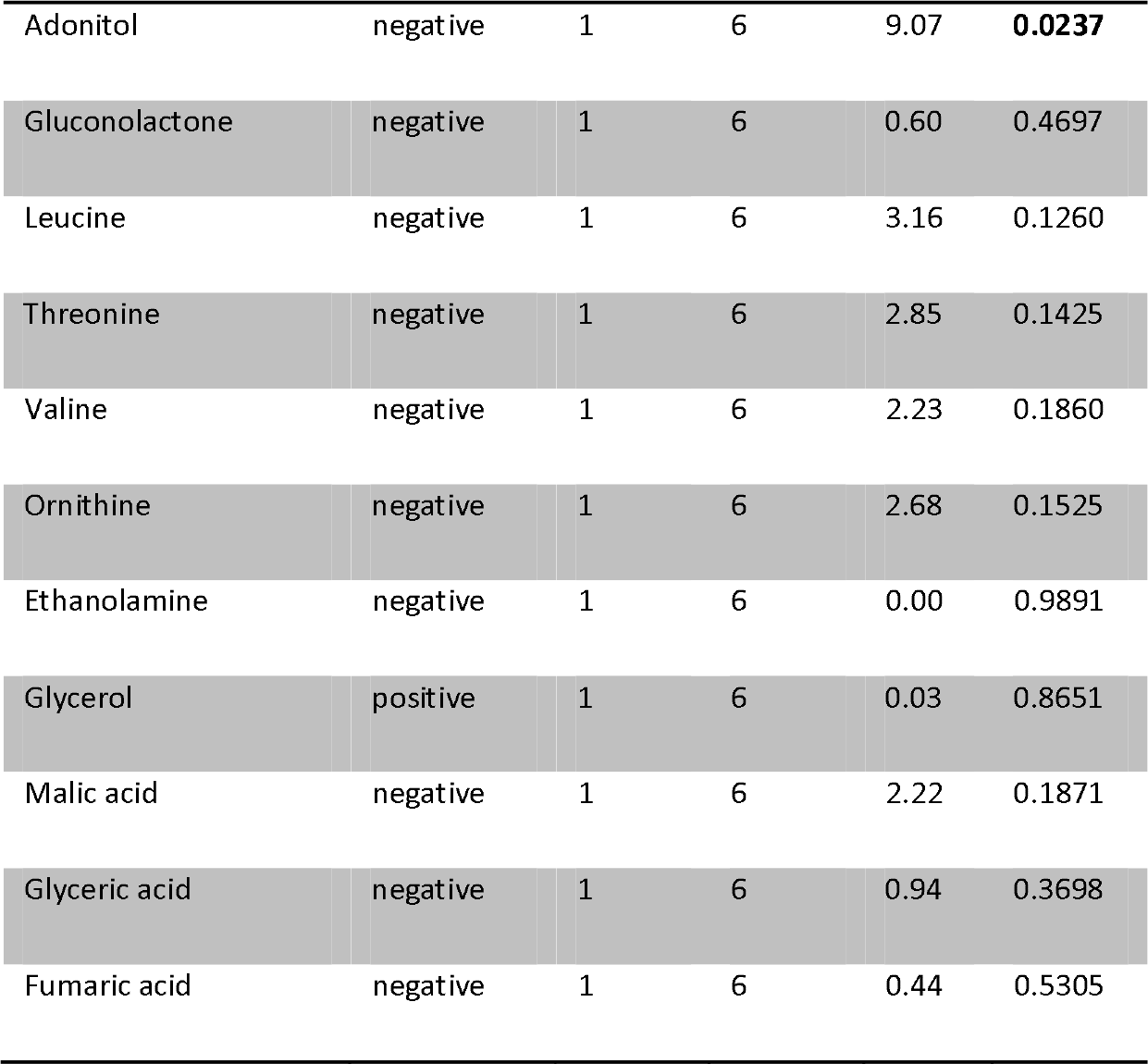

**E.**
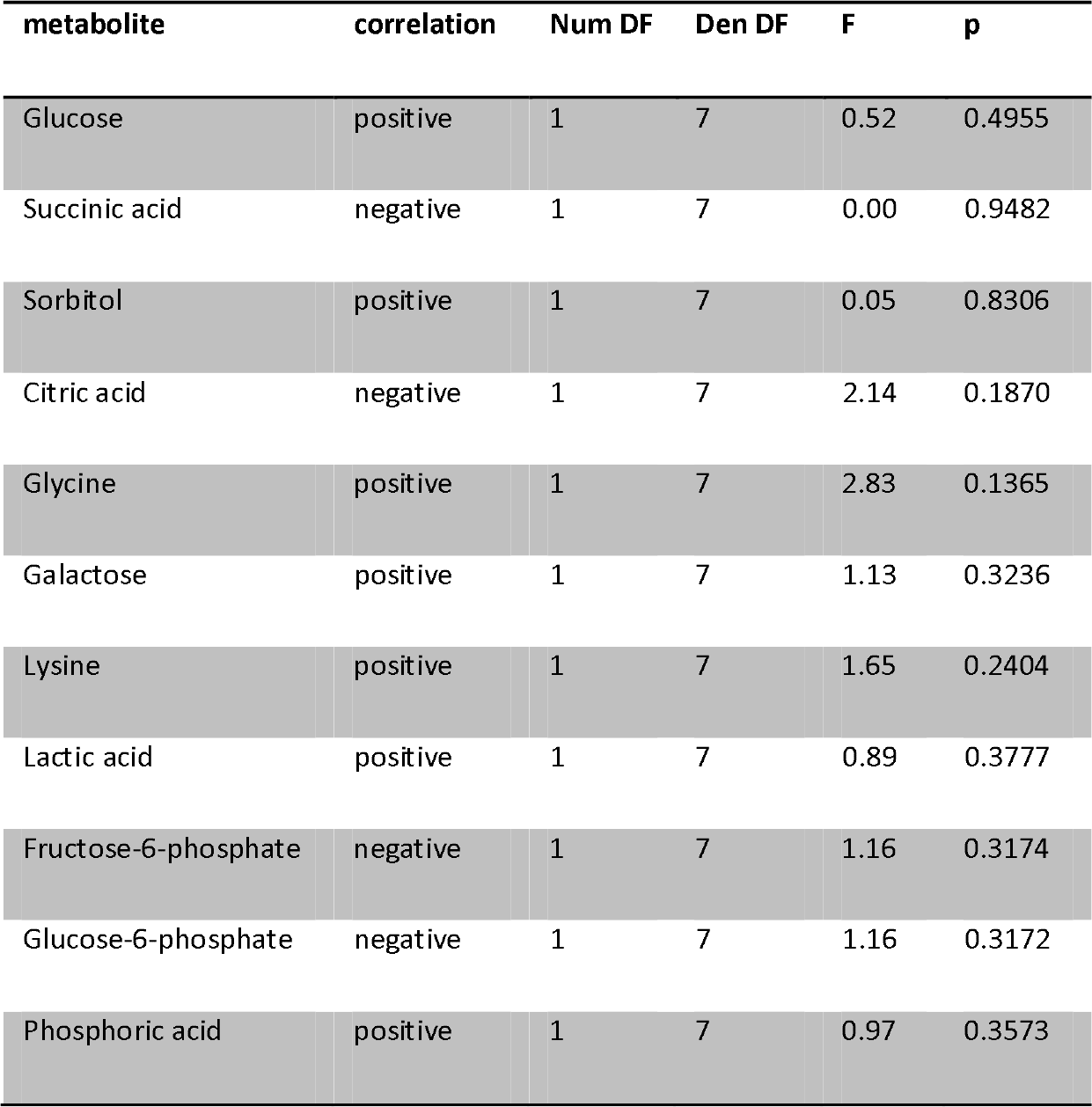
longevity

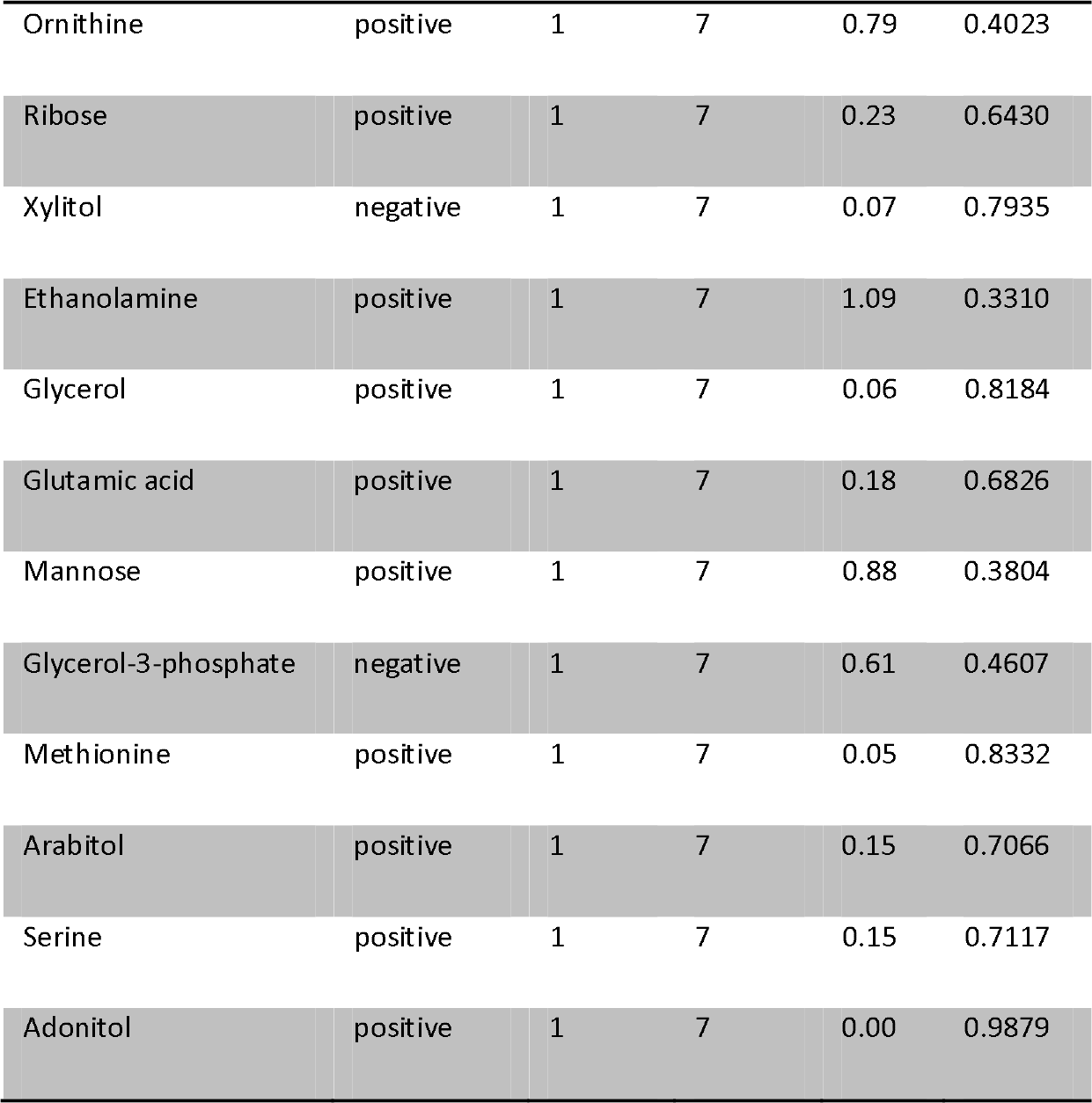

**F.**
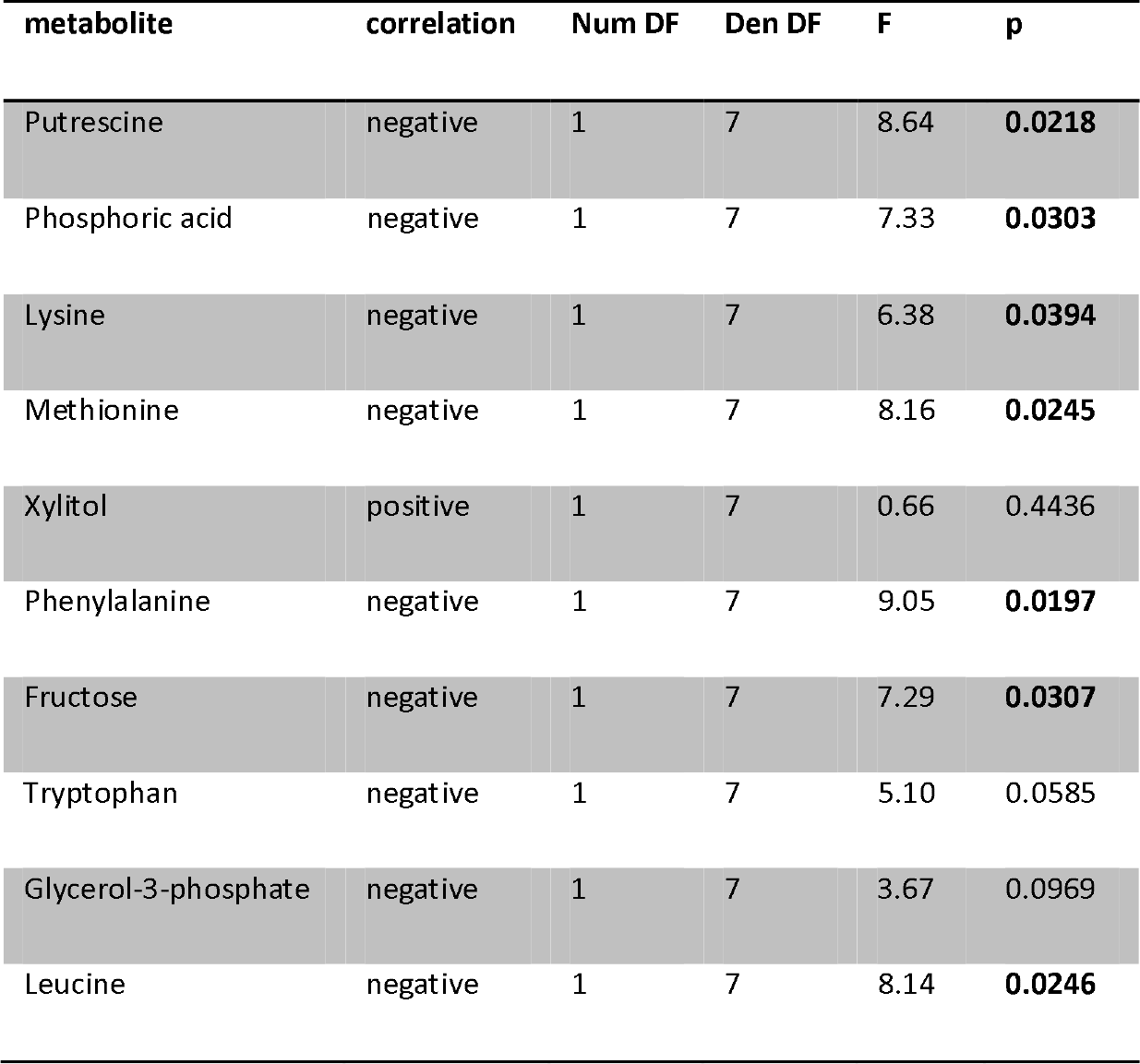
dispersal propensity

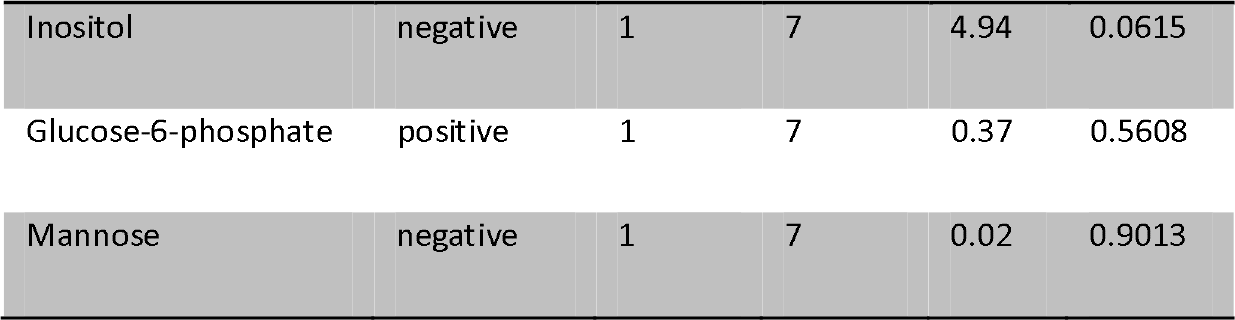

**G.**
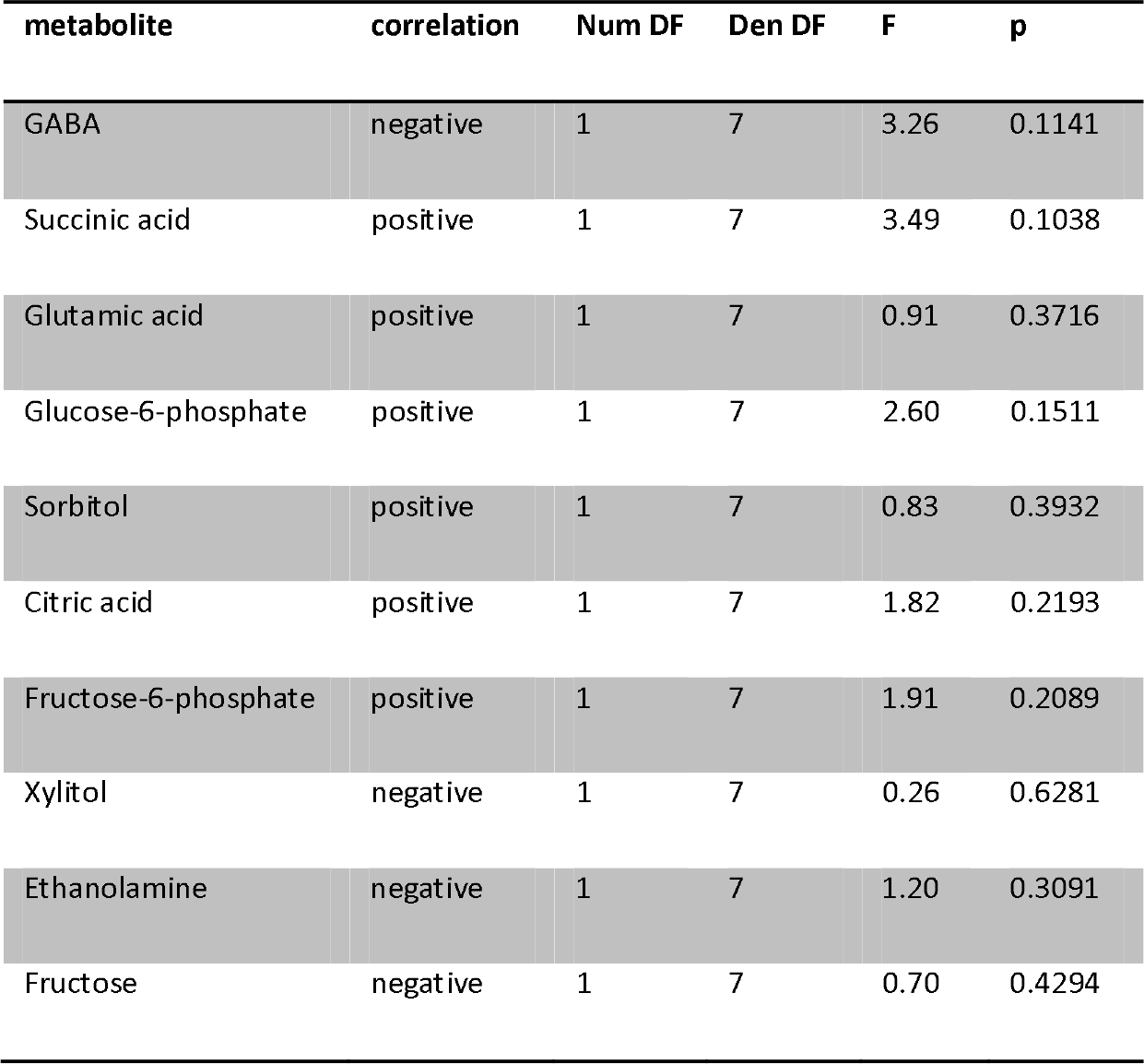
sex ratio

**G. sex ratio**
| metabolite | correlation | Num DF | Den DF | F | p |
| --- | --- | --- | --- | --- | --- |
| GABA | negative | 1 | 7 | 3.26 | 0.1141 |
| Succinic acid | positive | 1 | 7 | 3.49 | 0.1038 |
| Glutamic acid | positive | 1 | 7 | 0.91 | 0.3716 |
| Glucose-6-phosphate | positive | 1 | 7 | 2.60 | 0.1511 |
| Sorbitol | positive | 1 | 7 | 0.83 | 0.3932 |
| Citric acid | positive | 1 | 7 | 1.82 | 0.2193 |
| Fructose-6-phosphate | positive | 1 | 7 | 1.91 | 0.2089 |
| Xylitol | negative | 1 | 7 | 0.26 | 0.6281 |
| Ethanolamine | negative | 1 | 7 | 1.20 | 0.3091 |
| Fructose | negative | 1 | 7 | 0.70 | 0.4294 |

## A.7 Pathway Maps

Pathway maps of the two pathways in which some of the metabolites identified in the GC-MS analysis (visualised with either red or blue rectangles) play an important role. In the aminoacyl-tRNA biosynthesis pathway, the metabolites in the red rectangles correlate negatively with latitude and the blue ones correlate negatively with dispersal propensity. In the valine, leucine and isoleucine biosynthesis pathway, the metabolites in the red rectangles correlate negatively with latitude. The maps were used from KEGG (Kyoto Encyclopedia of Genes and Genomes) (Kanehisa *et al.* 2015).

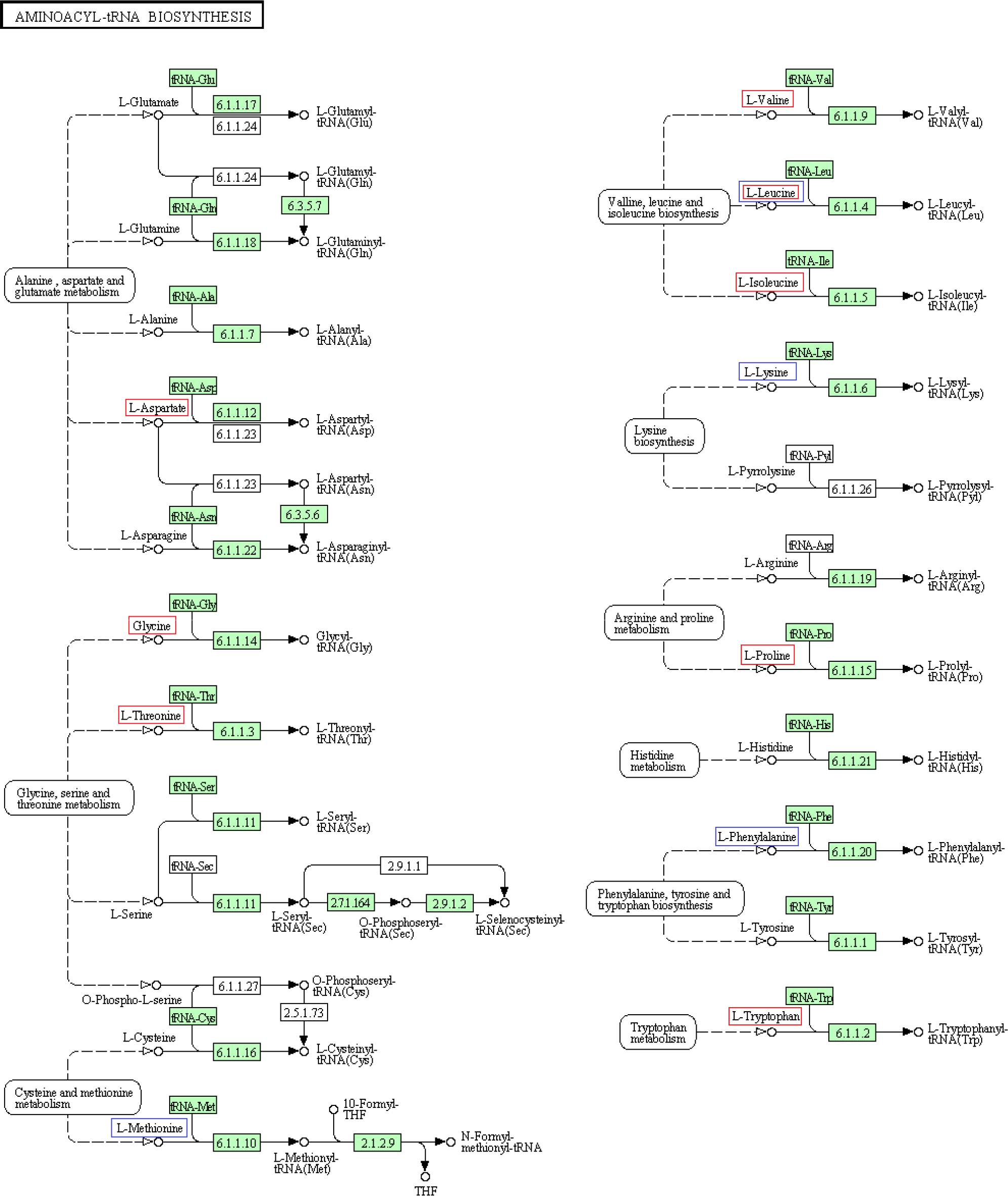

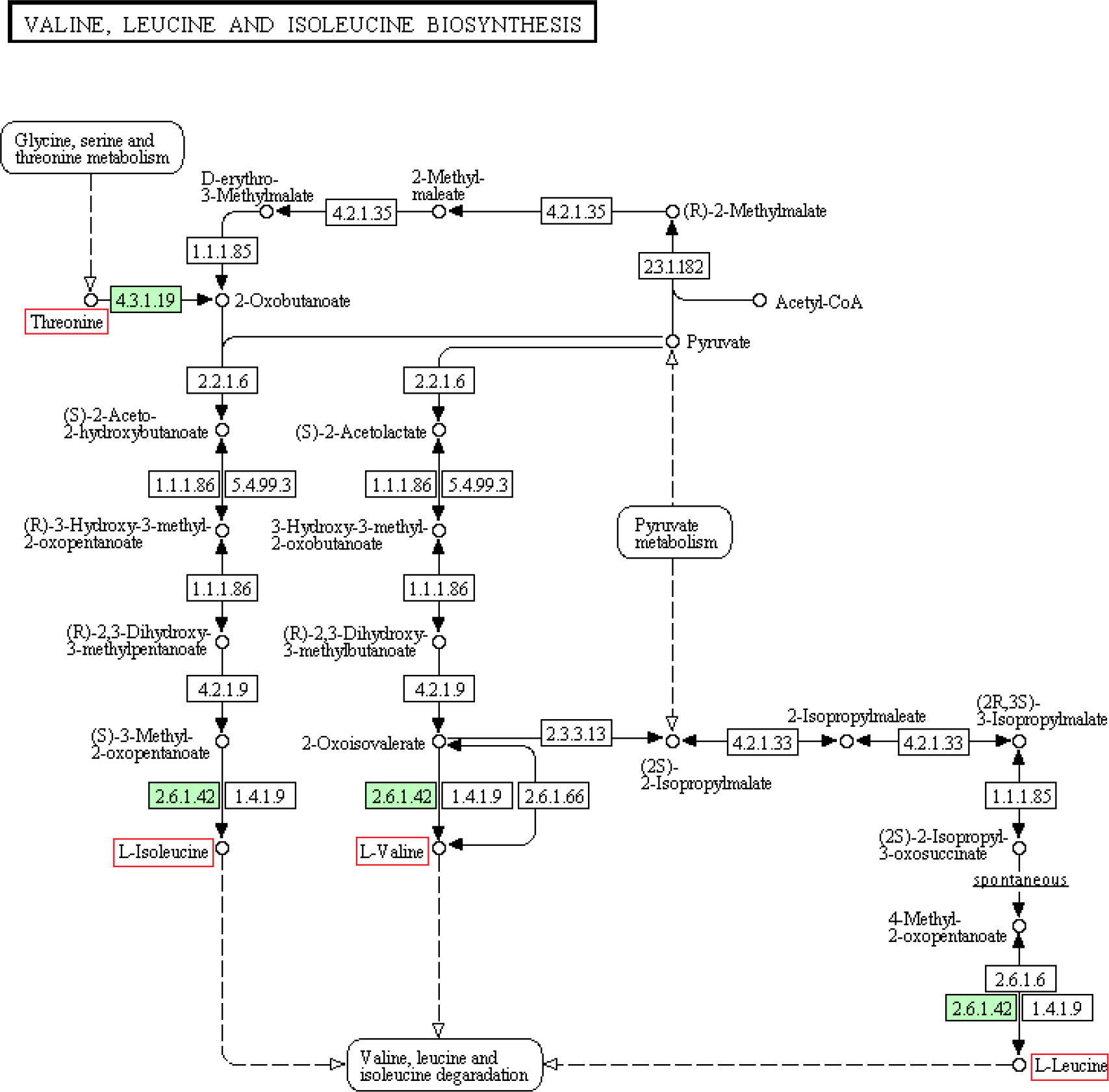

## A.8 Regression for all Populations *Versus* for *L. Periclymenum* Only

For those metabolites that gave significant results (see appendix A.6) for latitude or a specific life-history trait (daily fecundity, lifetime fecundity, egg survival, longevity, dispersal propensity or sex ratio) in the regressions run for all nine populations (hence for *L. periclymenum*, *Euonymus europaeus*, *Sambucus nigra* and *Humulus lupulus*), we repeated these regressions but then for *L. periclymenum* only (hence for only six populations-see appendix A.1). The table gives the slopes and p-values of both the regressions run for all populations (slope_all and p_all) and those run for *L. periclymenum* only (slope_*per* and p_*per*).

| trait | metabolite | slope_per | slope_all | p_per | p_all |
| --- | --- | --- | --- | --- | --- |
| latitude | Proline | -0.04354 | -0.02733 | 0.0205 | 0.0164 |
| latitude | Tryptophan | -0.05170 | -0.04474 | 0.1766 | 0.0383 |
| latitude | Inositol | -0.05628 | -0.04890 | 0.1880 | 0.0389 |
| latitude | Aspartic acid | -0.01956 | -0.02840 | 0.2757 | 0.0369 |
| latitude | Glycine | -0.00681 | -0.03254 | 0.6500 | 0.0315 |
| latitude | Malic acid | -0.00703 | -0.03858 | 0.6992 | 0.0372 |
| latitude | Valine | -0.02914 | -0.02727 | 0.1726 | 0.0245 |
| latitude | Isoleucine | -0.03947 | -0.02997 | 0.0947 | 0.0209 |
| latitude | Quinic acid | 0.02094 | 0.01594 | 0.2119 | 0.0500 |
| latitude | Threonine | -0.02807 | -0.02700 | 0.2046 | 0.0301 |
| latitude | Leucine | -0.01727 | -0.02270 | 0.3466 | 0.0359 |
| egg survival | Adonitol | -0.00651 | -0.00258 | 0.0751 | 0.0237 |
| dispersal propensity | Putrescine | -0.00092 | -0.00108 | 0.0854 | 0.0218 |
| dispersal propensity | Phosphoric acid | -0.10219 | -0.13336 | 0.0492 | 0.0303 |
| dispersal propensity | Lysine | -0.00277 | -0.00553 | 0.0942 | 0.0394 |
| dispersal propensity | Methionine | -0.00793 | -0.01081 | 0.1461 | 0.0245 |
| dispersal propensity | Phenylalanine | -0.01652 | -0.02201 | 0.1208 | 0.0197 |
| dispersal propensity | Fructose | -0.00266 | -0.00221 | 0.0744 | 0.0307 |
| dispersal propensity | Leucine | -0.03211 | -0.04462 | 0.1224 | 0.0246 |

